# Climate variability disrupts mutualism-driven increases in population persistence

**DOI:** 10.1101/2025.09.04.674083

**Authors:** Vicki W. Li, Joshua C. Fowler, Aaron S. David, Sharon Y. Strauss, Christopher A. Searcy, Michelle E. Afkhami

## Abstract

Understanding how species interactions impact population dynamics and long-term persistence over broad temporal and spatial scales is crucial for predicting species distributions and responses to global change. Here, we integrate range-wide field surveys of ∼90 grass host populations spanning 13 years with demographic modeling based on six-year common garden experiments conducted across the host range to demonstrate that mutualistic fungal endophytes promote population-level persistence and growth of their native host grass across its distribution, with non-mutualistic populations four times more likely to go locally extinct. Despite providing population-level benefits, endophyte prevalence declined eight-fold more in historically mutualistic populations that experienced high climate variability. Thus, mutualisms can underpin population persistence and buffer hosts against environmental stress, but may themselves be vulnerable to global change, with concerning implications for long-term population viability and, ultimately, species distributions under an increasingly uncertain climate.

## Introduction

Under global change, many species are experiencing dramatic range shifts and local extinctions^1–3^. In order to predict long-term species viability, it is crucial to understand what drives the persistence of individual populations across large geographic scales. One potentially important determinant of population persistence is mutualism, or positive interspecific interactions, wherein partners provide reciprocal fitness benefits. Because mutualisms can ameliorate the impacts of both biotic and abiotic stressors^4,5^, they may be especially important in the Anthropocene by enabling the persistence of organisms experiencing global change stress. However, empirical research scaling up the effects of mutualism to population persistence, especially across large spatial scales, has thus far been limited, with most studies focusing either on individual-level fitness^6,7^, local-scale population outcomes^8^, or correlated declines in occurrence records of mutualistic partners^9^. In this study, we employ a multifaceted approach directly linking mutualism to long-term persistence and extinction dynamics of natural populations. We bring together multiple lines of evidence, including surveys of host plant populations and the prevalence of microbial mutualists, large-scale field experiments, and demographic modeling, to connect the mechanistic effects of mutualism on population dynamics to real-world population outcomes in the context of increasing environmental stress and variability under global change. By doing so, we address two key questions. First, do microbial mutualisms promote population persistence and reduce extinction risk at large spatial and temporal scales, particularly in stressful or variable environments? And second, which environmental factors promote or disrupt the prevalence of these crucial mutualistic relationships? Answering these questions will help us understand and predict how species respond to global change.

Population persistence—a key component of both species viability and distributions—can be negatively impacted by abiotic forces such as drought and warming that are increasing in the Anthropocene^2,10^, potentially leading to local population extinctions. In contrast, beneficial mutualistic interactions can positively affect population dynamics by ameliorating stressors such as drought and may thus enable population persistence and even range expansions under abiotic stress^11–13^. Mutualisms may also ameliorate complex aspects of global change, including climate variability and extreme events such as fire, which could in turn scale up to promote population growth and range expansion^14,15^. However, how these complex, long-term aspects of environmental stress such as climate variability and extreme events shape mutualism effects on populations is not well-understood, especially in natural ecosystems.

Although mutualisms could be crucial for population persistence under increasing stress in the Anthropocene, they may themselves be disrupted by global change. The population-level benefits conferred by a mutualistic interaction are in part a product of its prevalence, or the frequency with which it occurs^16–19^. Therefore, over time, declines in mutualism prevalence could impact population persistence, species ranges, and, ultimately, species viability. Because mutualism benefits can vary with factors such as aridity^13,20^, the prevalence of mutualisms can depend on ecological context. For instance, prevalence of mutualistic fungal endophytes is often higher under more arid conditions^21^ and greater herbivory pressure^22^, reflecting how they benefit hosts by ameliorating drought or herbivory stress^23–25^. Overall, mutualisms are expected to persist when the benefits they confer exceed the costs of maintaining them and decline in prevalence otherwise^26^. However, environmental change, both over time and space, could affect mutualism prevalence by shifting the balance of costs and benefits. In particular, global change could increase reliance on stress ameliorating mutualisms or, alternatively, disrupt mutualistic interactions^27^. Global change could also reduce mutualism prevalence through means that are independent of costs and benefits, such as through creating phenological mismatches between partners^28^. Taken together, this emphasizes the importance of simultaneously tracking mutualism impacts on population outcomes and global change impacts on mutualism prevalence. Determining which environmental factors promote or disrupt the prevalence of mutualisms is crucial to predicting whether these beneficial interactions will continue to occur in a changing world and, in turn, underlie long-term population persistence.

Here, we explore how mutualism scales up to impact long-term population dynamics across species ranges, especially under environmental stress, and how global change has influenced the prevalence of these crucial mutualistic relationships. To do so, we perform 1) field surveys of ∼90 widely distributed natural host populations 13 years apart to identify differential persistence of mutualistic and non-mutualistic populations as well as changes in mutualism prevalence, 2) two sets of six-year common garden experiments spanning 92% of the host’s climatic range to determine mutualist effects on host vital rates across space and time, and 3) range-wide demographic modeling of mutualist-associated and non-associated populations to uncover the pathways through which mutualisms affect population trajectories. We conduct this research using the mutualism between the California-native grass *Bromus laevipes* and its symbiotic systemic foliar endophytic fungi^29^. In general, endophytic fungi are ubiquitous and occur in every major plant lineage^30^, with vertically transmitted systemic fungal endophytes of the genus *Epichloë* (Clavicipitaceae) occurring in up to ∼40% of species across cool-season grasses^31–34^. These widespread symbionts have also been shown to expand their host species range by conferring drought tolerance, although they may be costly to support in wetter habitats^13^. The endophytic fungi’s established ability to ameliorate drought stress in our model grass species, combined with its natural variation in prevalence and a growing recognition of the importance and ubiquity of aboveground microbes, make this system ideal for investigating how microbial mutualists influence plant population dynamics under global change.

Overall, we hypothesized that these microbial mutualists would provide range-wide benefits to host populations, especially in stressful or variable environments. We further hypothesized that mutualism prevalence would increase under those conditions where it was most beneficial. Our results revealed for the first time that microbial mutualisms promote population growth and persistence of their plant host across its distribution, yet climate variability—a key component of global change^35^—reduces mutualism prevalence, undermining the very interactions that support host resilience. These results uncover a critical paradox—that mutualisms that buffer plant populations against environmental stress may themselves be vulnerable to the destabilizing forces of global change.

## Results

### Local extinction over 13 years was four times higher in non-mutualistic plant populations

To understand the importance of endophyte mutualism to host plant population persistence, we surveyed 86 natural populations of *B. laevipes* across its range in northern and central California in 2009–10 and again 13 years later in 2022, evaluating persistence as well as endophyte prevalence at both time points for each population (>2700 host plants evaluated; Figure 1; see Methods). After determining that ∼27% of populations went locally extinct, we tested how host persistence across the 13 years and current endophyte prevalence (in 2022) of each population depended on historical endophyte prevalence (in 2009) and its interactions with environmental variables relevant to global change (mean climate, change in mean climate, climate variability, and fire; climate was quantified using SPEI, a drought/aridity index accounting for temperature and precipitation, with lower values indicating greater drought^36^). Global model selection on candidate multiple logistic regression models identified that the best supported persistence model included two factors—historical endophyte prevalence and fire between surveys—important for predicting *B. laevipes* population persistence (P = 0.012), with no interaction effect between the two (Z = 1.45, P = 0.15). Interestingly, as historical endophyte prevalence within populations increased, populations were significantly more likely to persist (χ1^2^ = 6.73, P = 0.0095; Figure 2). Populations with no endophytes detected in 2009-2010 were roughly four times more likely to go locally extinct than populations with fixed (i.e., 100%) endophyte mutualism in 2009-2010 (predicted probability of extinction: 45% for non-mutualistic populations vs 12% for fixed populations if fire occurred; 25% vs 5.1% if no fire occurred). We also found that plant populations were somewhat less likely to persist if fire had occurred between surveys (χ1^2^ = 2.91, P = 0.088; Figure 2). Overall, these results show that plant population persistence increased with endophyte prevalence, independent of whether fire has increased the probability of local extinction.

**Figure 1.**
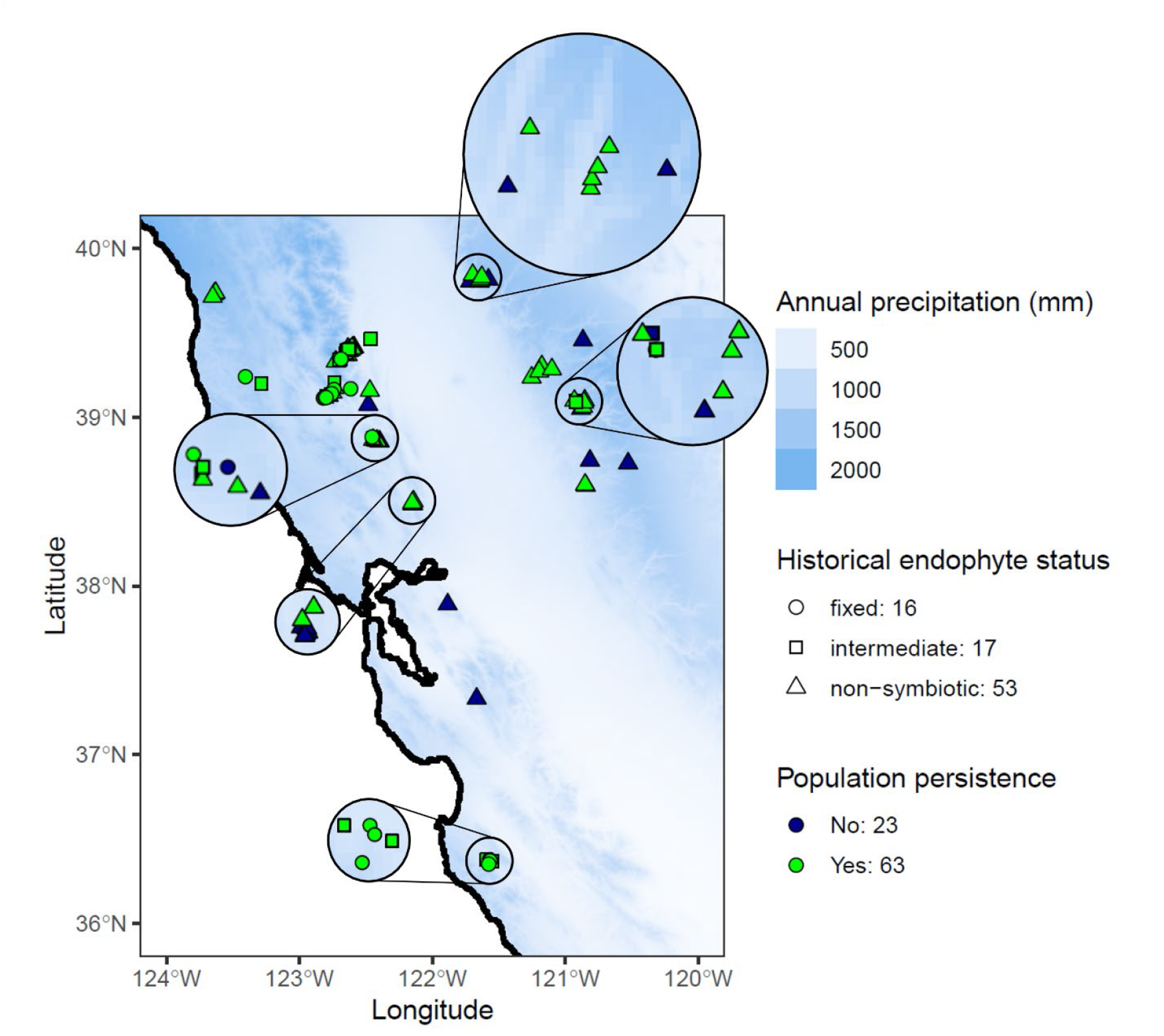
After 13 years, ∼27% of monitored plant populations went locally extinct, with populations varying in persistence within and across regions of northern and central California (which spans much of this host’s climatic range and geographic distribution; ∼98% of *B. laevipes* population records are within the California floristic province, with highest densities in northern and central California^13^). Map colors represent annual precipitation (mm; data from WorldClim). Each point represents one population surveyed in 2009–10 and in 2022. Each point’s color corresponds to whether or not it persisted to 2022. Each point’s shape corresponds to its historical (2009) endophyte status (fixed, prevalence > 90%; intermediate, 10% < prevalence < 90%; non-mutualistic, prevalence < 10%; cutoffs based on previous work^13^). Numbers following legend labels indicate how many populations fall under each category (e.g., 63 populations persisted to resampling). (Also see Table S2 and Figure S4.) Note that *B. laevipes* does not occur in the Central Valley (roughly 40°N, 123°W to 37°N, 120°W), which is dominated largely by invasive annual grasses, according to both our surveys and herbarium records.

**Figure 2.**
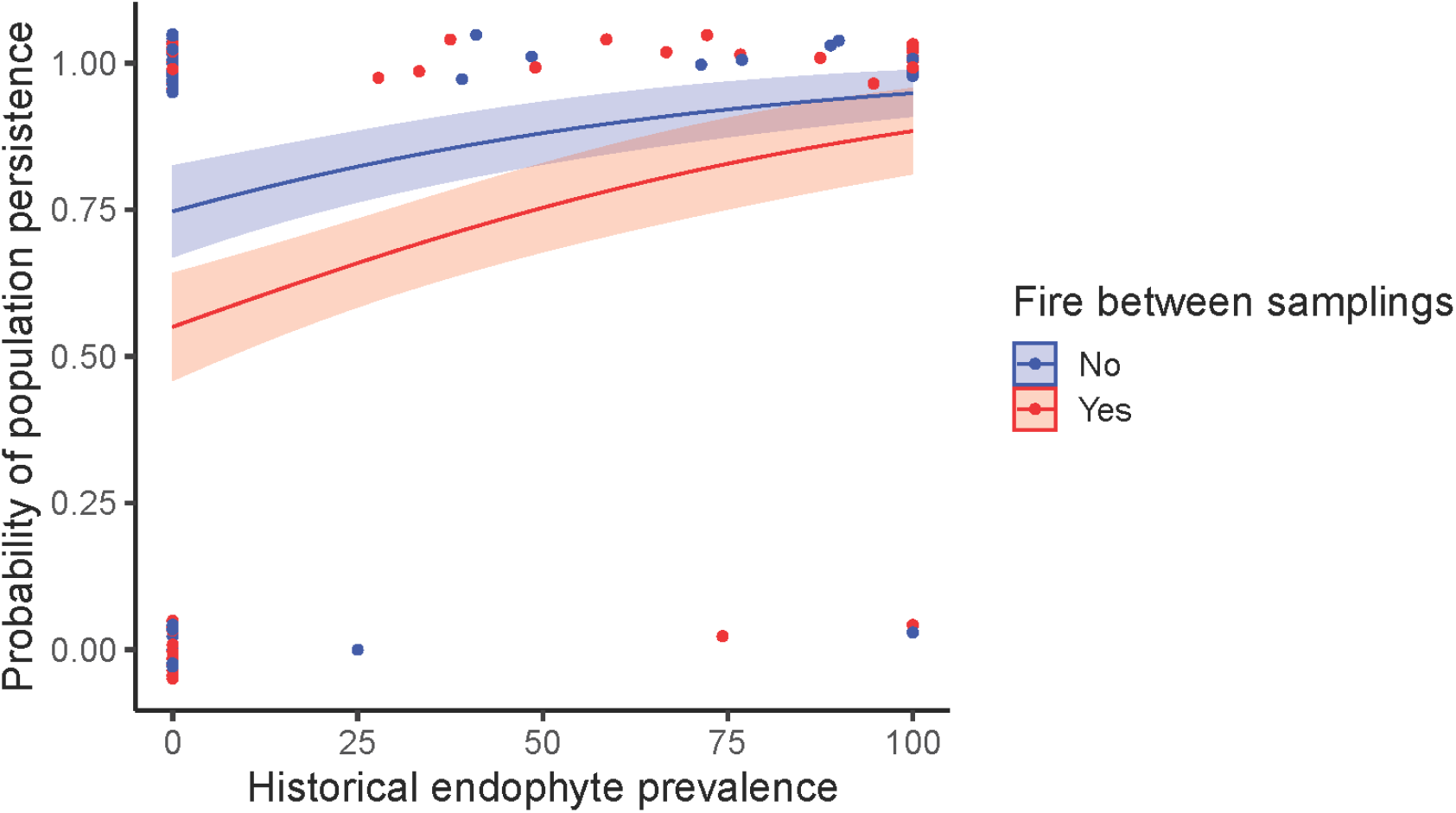
Plant population persistence was explained by historical endophyte prevalence (χ1^2^ = 6.73, P = 0.0095) and the occurrence of fire between surveys (χ1^2^ = 2.91, P = 0.088) in our best supported statistical model, with probability of population persistence increasing with historical endophyte prevalence and decreasing with fire occurrence. Historical endophyte prevalence refers to the endophyte prevalence of a *B. laevipes* population in 2009 during the initial field surveys. Each point represents one *B. laevipes* population, with color corresponding to whether or not fire had occurred between surveys. Points have been jittered for visualization purposes. The model is a logistic regression and the shaded areas represent the standard error. Figure created using *ggPredict*^37^.

### Mutualistic host populations had ∼30% higher population growth rates driven by endophyte-enhanced fecundity

To assess the effect of endophyte mutualism on plant population dynamics, we performed two sets of common garden experiments involving 1650 plants and 550 seeds, monitored for six years at five sites chosen to span the *B. laevipes* range and a wide ecological and climate gradient (northern to central California; ∼420 km; ∼450–1750 mm average annual precipitation during experiment years). We then measured and parameterized vital rates and constructed population projection matrix models to test how predicted finite population growth rates (λ) depended on endophyte mutualism, as well as performed fixed-effect life table response experiments to assess whether endophytes impacted population dynamics through endophyte effects on host growth and survival or via host fecundity. Using our demographic model, we found that λ differed significantly between sampled populations, with endophyte-associated populations having 28.37% higher population growth rates on average than endophyte-free populations (pseudo-P < 0.0001; λ = 1.67 ± 0.021 for endophyte-associated populations vs λ = 1.30 ± 0.025 for endophyte-free populations; Figure 3). Furthermore, our life table response experiment revealed that these endophyte benefits to population growth originated primarily through impacts on fecundity rather than on growth and survival (Wilcoxon signed-rank test: V = 3741, Z = 8.05, P < 0.0001; contribution: -0.045 ± 0.0055 through survival/growth and 0.35 ± 0.0058 through fecundity, with the difference between them being 0.39 ± 0.011; mean ± SE; Figure 3). This life table response experiment was performed on demographic model predictions for each surveyed population to generalize across *B. laevipes*’s species range. Specifically, we tested how much endophyte association was predicted to increase λ for each population and partitioned that contribution to population growth between life class transitions and between survival/growth and fecundity.

**Figure 3.**
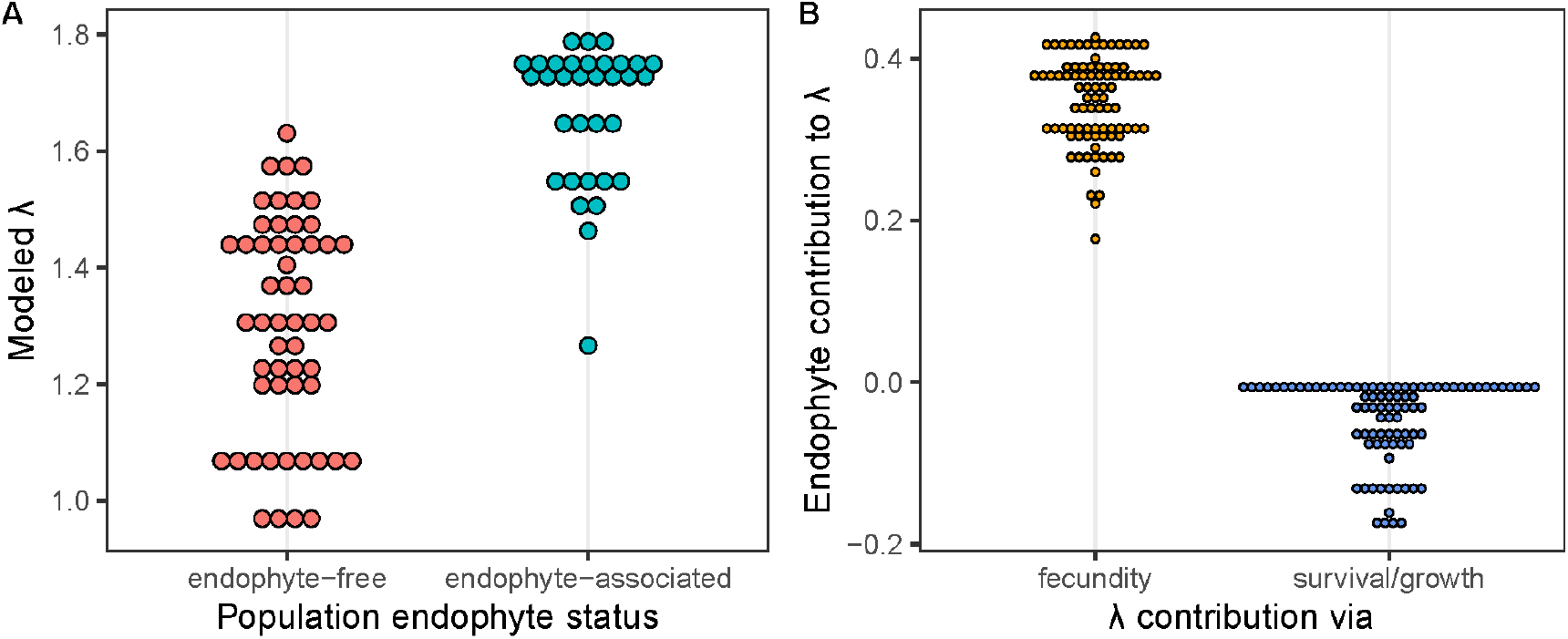
**(A)** Population growth rates of endophyte-associated populations were significantly greater (pseudo-P < 0.0001) than those of endophyte-free populations based on population growth rates projected for each of the 86 surveyed populations across the *B. laevipes* range (53 endophyte-free, 33 endophyte-associated). Endophyte-associated populations include both intermediate and 100% endophyte prevalence (fixed) populations. **(B)** Modeled endophyte mutualism contributions to population growth rates were greater through fecundity than through survival/growth (V = 3741, Z = 8.05, P < 0.0001). Here, λ contribution refers to how much our model predicted endophytes would increase λ for each population (i.e., predicted λ when endophyte-associated minus predicted λ when endophyte-free). Projections are based on a demographic model parameterized with data from our multi-site, multi-year common garden experiments. Also see Figures S10 and S11.

### Endophytes underlie host persistence in arid climates, but endophyte benefits decline under more variable climates

Using our demographic model, we calculated endophyte contribution to population growth rate (i.e., how much endophytes would increase population growth rate) as the difference between predicted endophyte-associated and endophyte-free λ values for each of the 86 surveyed populations spanning the *B. laevipes* range. We found that broadly, as mean SPEI values across the four experimental years increased (i.e., as sites became wetter), predicted λ values increased for both endophyte-associated and endophyte-free populations (Figure 4). Notably, in the driest of the sites occupied by *B. laevipes* populations, endophyte association allowed hosts to sustain positive population growth rates (i.e., λ > 1) that were not achieved by endophyte-free populations, suggesting that in extremely dry conditions, endophyte mutualism is required for population persistence.

**Figure 4.**
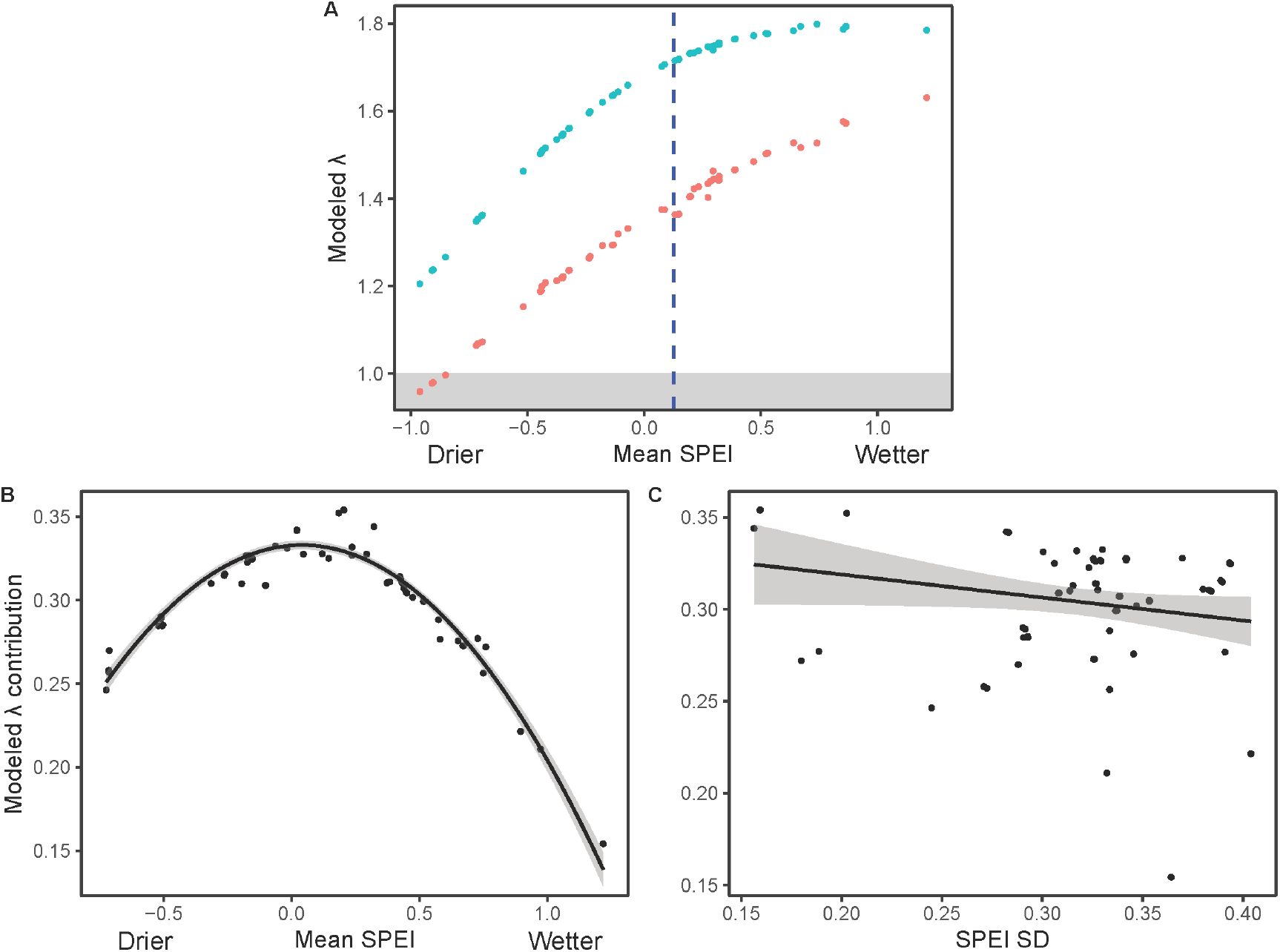
**(A)** Modeled population growth rates, both endophyte-free and endophyte-associated, for the 86 sites corresponding to our field survey populations across the *B. laevipes* range. As sites became wetter on average during the four experimental years, predicted λ values increased for both endophyte-associated and endophyte-free populations. Endophyte contributions to modeled population growth rates for the 86 sites (i.e., the difference in modeled λ between endophyte-associated and endophyte-free populations) **(B)** had a unimodal relationship with aridity (R^2^ = 0.93, F_2,83_ = 584.3, P < 0.0001) and **(C)** decreased marginally significantly with variability in aridity (R^2^ = 0.042, F_1,84_ = 3.68, P = 0.058), where each point represents one *B. laevipes* population, the line is the polynomial or linear regression, and the shaded areas represent the standard error. Here, mean SPEI refers to the SPEI of each site in the *B. laevipes* range, averaged across the years of the common garden experiments. The dashed blue line represents the mean SPEI for all 86 surveyed sites of *B. laevipes* populations over the four experimental years; note that the mean is not centered at zero because the climate values from the four common garden experimental years represent only a subset of the climate data used to calculate SPEI values. SPEI SD refers to the standard deviation of SPEI of each given site over the same years. Note that vital rate functions were parameterized using SPEI as a candidate term, which is in turn closely related to the mean SPEI across experimental years at each site. Thus, for Figure 4B in particular, model predictions are strongly explained by site mean SPEI, with deviations from the unimodal curve largely owing to year-to-year climate variability. Figure 4B therefore primarily illustrates that model predictions do indeed follow a unimodal curve.

The endophyte contribution to population growth rate was positive across all sites, indicating that endophyte mutualism provided a range-wide demographic advantage to its host. Importantly, relative endophyte benefits to population growth rate had a unimodal relationship with mean SPEI across the experimental years, with the greatest endophyte contributions to population growth occurring in sites with intermediate to slightly dry conditions (R^2^ = 0.93, F_2,83_ = 584.3, P < 0.0001; Figure 4). Furthermore, modeled endophyte contributions to population growth rates declined with increasing climatic variability across the experimental years (R^2^ = 0.042, F_1,84_ = 3.68, P = 0.058; Figure 4). Overall, because of the difference in climatic optima for endophyte-associated and endophyte-free populations, the relative benefits of endophyte mutualism were greatest in roughly average conditions and therefore declined with increasing climate variability (see Supplementary Discussion: *The contribution of endophyte mutualism to population growth declines with increasing climatic variability*).

### Endophyte prevalence declined in historically mutualistic host populations under high climate variability

Although results from both surveys of natural populations and demographic modeling indicate endophytes were broadly beneficial, endophyte prevalence did not increase over time in surveyed field populations (see Supplementary Discussion: *Host populations maintained intra-population variation in mutualism rather than progressing towards endophyte fixation*). Thus, to understand the relationship between current and historical endophyte prevalence among all surveyed populations and what environmental factors promote or undermine endophyte prevalence, we conducted global model selection using the same candidate explanatory terms as our population persistence model (see Methods). Three factors—historical endophyte prevalence (χ1^2^ = 27.35, P < 0.0001), variability in aridity (the standard deviation of SPEI; χ1^2^ = 10.52, P = 0.0012), and aridity (mean SPEI; χ1^2^ = 3.31, P = 0.069)—were important for current endophyte prevalence (P < 0.0001; Figure 5). The interaction effect of the two significant explanatory terms, historical endophyte prevalence and standard deviation of SPEI, was also significant (Z = -2.93, P = 0.0017; see Supplementary Methods: *Historical endophyte prevalence, climate variability, and their interaction are important to current endophyte prevalence*). As expected, we found that historical and current endophyte prevalence are strongly related (i.e., populations with historically higher endophyte prevalence continue to have high endophyte prevalence and vice versa). Interestingly, under high climatic variability, historically highly mutualistic populations (90% endophyte prevalence) were predicted to lose their mutualists ∼8x more on average compared to populations with historically low (10%) mutualism (historically high mutualism: 67.1% loss in prevalence versus historically low mutualism: 7.9% loss). Furthermore, predicted loss of mutualists from high mutualism populations under high climatic variability (67.1% loss) was far greater than gain of mutualists in low mutualism populations under low climatic variability (15.1% gain), suggesting differential responsiveness of endophyte prevalence to climate variability for high and low mutualism populations.

**Figure 5.**
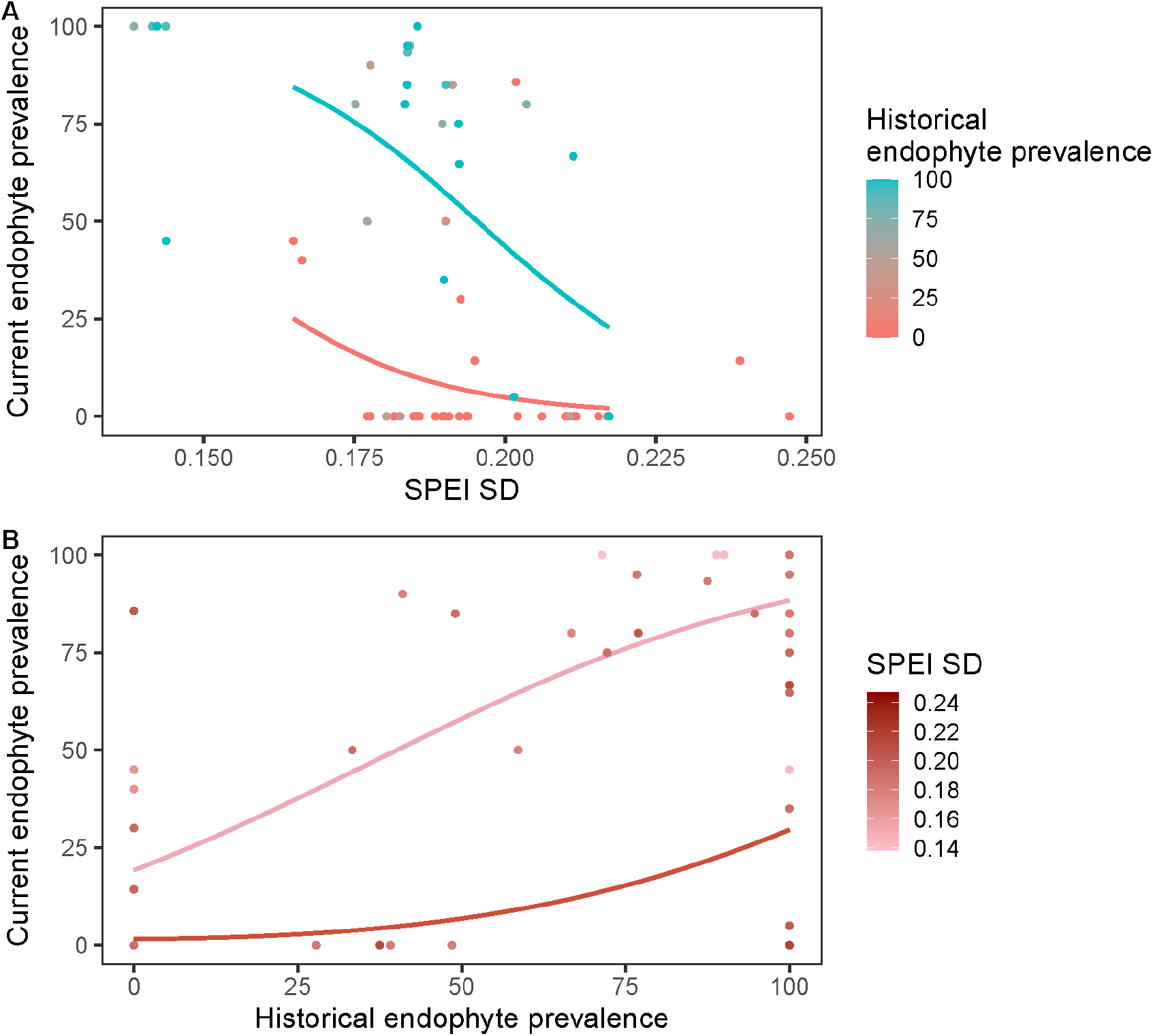
Current endophyte prevalence in 2022 resurveys plotted against **(A)** the standard deviation of SPEI and **(B)** historical endophyte prevalence in the original 2009 surveys. Current endophyte prevalence was generally positively related to historical endophyte prevalence, but endophyte prevalence has declined over time in historically highly mutualistic populations with higher climate variability. Points represent *B. laevipes* populations, and their colors represent their **(A)** historical endophyte prevalence and **(B)** climate variability experienced. The curves are the regression model plotted for **(A)** both high (90%) and low (10%) levels of historical endophyte prevalence and **(B)** both high (0.22) and low (0.16) levels of climate variability. The regression model shows that current endophyte prevalence is a function of historical endophyte prevalence (χ1^2^ = 27.35, P < 0.0001), interannual variation in aridity (i.e., standard deviation of SPEI; χ1^2^ = 10.52, P = 0.0012), and mean aridity (χ1^2^ = 3.31, P = 0.069); the interaction effect between current endophyte prevalence and variation in aridity was also significant (Z = -2.93, P = 0.0017).

## Discussion

By combining large-scale common garden field experiments, demographic modeling, climate data, and long-term population monitoring across the host species’ range, our research provides two novel insights. First, as we hypothesized, microbial mutualists can underpin long-term plant population persistence across species ranges. Our demographic model predicted that facultative mutualistic microbes enhance host population viability across the range, in line with significantly greater rates of persistence of mutualistic natural populations over 13 years from our surveys. Second, variability in a climate change-related stress (i.e., aridity) can select against a mutualism that ameliorates directional changes in the mean of the same stressor. Despite endophyte mutualism being broadly beneficial to host populations in this study, its prevalence did not increase within natural populations and endophyte loss was highest among populations experiencing greater temporal climate variability, potentially undermining the very interactions that support hosts in arid conditions. Our population model additionally revealed that endophytes promote host population growth primarily through enhancing fecundity. Furthermore, our demographic model showed endophytes can underpin persistence in sites with the driest climates, which scales up our previous research showing endophytes confer drought tolerance to individual plants and allow hosts to occupy the driest parts of their range^13^. However, we found no evidence that endophytes buffer hosts from climate variability or extreme events, nor did we observe increases in endophyte prevalence, which was contrary to our second hypothesis. In fact, results from both our demographic predictions and surveys of natural populations support that more variable climates weaken endophyte benefits and reduce their prevalence, respectively, despite the broadly beneficial nature of endophyte mutualism.

### Mutualistic endophytes support host population persistence and growth

Our field surveys and demographic model showed endophyte mutualism positively impacts range-wide host population persistence and growth rate, respectively. Notably, previous demographic modeling approaches have also predicted that microbial mutualisms contribute to plant population growth^11,15,38–40^. However, our study is the first to complement predictions of increased population viability made from demographic modeling with monitoring of natural populations to show in real world field conditions that populations with these microbial mutualists actually have significantly higher persistence across the species range. Surveyed historically non-mutualistic populations were roughly four times more likely to suffer local extinction than their highly mutualistic counterparts—especially remarkable given that *B. laevipes* is a perennial grass and fewer than 15 years elapsed between surveys. Further tracking of natural population sizes would allow an even more in-depth understanding of mutualism impacts on population dynamics and extinctions. Overall, the combination of the growing body of demographic modeling research predicting microbial mutualists increase population growth^11,15,38–40^ and our new work validating that mutualistic populations do, in fact, have greater persistence through time and across space suggests microbial mutualist impacts on population persistence are widespread. Furthermore, this study demonstrates that demographic model predictions based on experimental populations align with population outcomes in real communities. The life table response experiment revealed that biologically, endophytes provide these demographic benefits largely through enhancing host fecundity, in line with dynamics in other grass–endophyte systems^41,42^. While this mutualism was overall beneficial for host population growth, the negligible or even negative effects of endophytes on particular vital rates related to host growth and survival may hint at subtle partner conflict (discussed further in Supplementary Discussion: *Endophyte effects on host vital rates suggest possibilities for host– mutualist conflict*).

Importantly, the effects of mutualism on populations could ultimately impact host species ranges. By enhancing fitness and modifying niches through abiotic stress amelioration, mutualisms can expand the set of environmental conditions and thus geographic ranges species persist in^43^. Previous empirical research has demonstrated that low prevalence of mutualists outside a species’ established range is linked to reduced individual fitness, highlighting how mutualisms can help set range limits^6,17,19,44–46^. Here, we demonstrated that low mutualist prevalence can also explain local extirpations *within* species ranges, a finding that, if not compensated for by dispersal, could manifest in range contractions over long timescales. Our demographic model predictions also provide ecologically mechanistic insight into how microbial mutualists can expand species ranges through abiotic stress amelioration. Endophyte mutualisms can enhance host tolerance to drought^25^, and our original study^13^ demonstrated how endophyte-conferred drought amelioration allows *B. laevipes* to expand its range into drier habitats. In support of this, our new demographic model showed endophyte association was necessary for host populations to achieve positive population growth rates in the driest parts of their range. Therefore, by ameliorating abiotic stress, endophyte mutualism enables *B. laevipes* populations to inhabit regions that are otherwise overly stressful^4^. In the context of global change, moisture deficit across *B. laevipes*’s range is projected to increase by >40% in some areas^47^. As mean climate becomes more arid, our demographic model predicts that endophytes may become indispensable to the persistence of *B. laevipes* and potentially other hosts.

### Climate variability selects against endophyte mutualism

Multiple lines of evidence implicated climate variability as a disrupter of endophyte mutualism. First, although our demographic model predicted consistently positive endophyte effects on population growth rates and that endophytes are crucial in highly arid conditions, endophyte contribution to population growth rate declined with increasing climate variability. Specifically, predicted endophyte contributions were 10.56% higher in the least variable site compared to the most variable one. This was due to endophyte-associated and non-associated populations having different climatic optima, leading to the *relative* benefit of endophyte mutualism to population growth from our demographic model being greatest in intermediate to slightly dry climates. Because variable climates by nature include more climatically extreme years, and in both directions (dry and wet) the relative benefit of endophyte mutualism is diminished, climate variability weakens positive selection for endophyte mutualism. Second, when surveying endophyte prevalence within natural host populations, we found that rather than trending upward, it in fact declined most strongly in populations with previously high mutualism prevalence and greater climate variability. Historical endophyte prevalence was also likely shaped by pre-survey (i.e., pre-2009) climate variability. However, climate variability has substantially increased across the vast majority of *B. laevipes*’s range in recent years, and has therefore likely only become more important in structuring mutualism among these populations (see Supplementary Discussion: *Climate variability structures endophyte prevalence*). Interestingly, while historically highly mutualistic populations declined in prevalence under high climatic variability, historically non-mutualistic populations did not see equivalent prevalence increases under low climatic variability. This asymmetry may result from mutualistic populations being more easily able to lose mutualists (e.g., via imperfect vertical transmission within mutualistic populations^48^) than non-mutualistic populations can gain mutualists (e.g., via migration of infected individuals into a population). Climate variability can impact mutualism prevalence in other systems (e.g., arbuscular mycorrhizal fungi colonization^49^). Importantly, our study revealed that variability in a climate change-related stress (i.e., aridity) can select against a microbial mutualism that ameliorates the same stressor. This negative selection occurred despite clear evidence that this mutualist increases host tolerance to drought in field and greenhouse experiments^13^ and, in this study, increases population growth above a critical threshold for maintaining persistence under severe aridity. Therefore, researchers should exercise caution when concluding that stress-ameliorating mutualisms should be selected for by global change, because mutualisms providing benefits under one parameter of a stressor (i.e., mean aridity) may simultaneously be weakened by another parameter in the same stressor (i.e., variability in aridity).

Variability selecting against mutualism was surprising. Given recent modeling research predicted endophytes provide demographic buffering that protects against environmental stochasticity^15^, we expected their prevalence to increase under more variable abiotic conditions. However, not only did we detect that endophyte prevalence declined in surveyed natural populations under high climate variability, but our demographic model also predicted decreasing benefits of endophyte mutualism with increasing climatic variability. Mutualisms can be environmentally context-dependent^50^, and in variable environments, where the costs and benefits of a microbial symbiont can change over time, models suggest the ability to lose microbial symbionts when they are costly improves host fitness.^51^ Taken together, this suggests climate variability reduces the benefits of endophytes relative to their costs, leading to declining prevalence. Climate variability could also select against endophyte prevalence through several additional pathways. First, climate variability could directly reduce transmission frequencies, which for vertically transmitted mutualists is linked to prevalence^52^. For instance, endophytes can confer thermal tolerance for their hosts, but higher temperatures have been associated with lower transmission frequencies in some grasses^7,53–56^. If endophyte-infected hosts are favored but endophyte transmission is reduced in hotter years that occur in more variable climates, endophyte effects could become unlinked from transmission and therefore prevalence. Alternatively, climate variability could promote coexistence between mutualistic and non-mutualistic individuals within populations if they exhibit niche differentiation^57^. Context-dependency in mutualisms can result in niche shifts and differentiation between mutualistic and non-mutualistic hosts, which temporal environmental variability can then act upon to promote coexistence of host types^57,58^. Previous research and our demographic modeling here suggest that endophyte-associated and endophyte-free *B. laevipes* populations occupy different climatic niches, with mutualistic populations occupying drier climates and vice versa^13^. If niche differences exist between mutualistic and non-mutualistic host individuals, a more heterogeneous environment through time may allow for coexistence, resulting in selection against mutualism in populations with relatively high prevalence and thus populations that are stable at intermediate rather than near-fixed levels of mutualism^59^. These scenarios demonstrate how a mutualism being broadly beneficial does not necessitate increases in its prevalence if it is impacted by climate variability.

### Conclusion and future directions

Our study demonstrates facultative microbial mutualisms can have outsized importance for the persistence of natural host plant populations across range-wide and decadal scales. However, the benefits of mutualism are not always accompanied by increases in mutualism prevalence; in fact, here, the prevalence of mutualism declined in populations experiencing more variable climates. Moving forward, many regions, including the range of *B. laevipes*, are anticipated to experience mean shifts in climate but also more variable climates^35,47^. Our findings demonstrate that even if mutualisms are generally beneficial or even favored by anticipated shifts in mean conditions— for example, towards drier climates—selection for mutualism prevalence to increase could be outweighed by increases in climate variability (e.g., in precipitation). This, unfortunately, may be ultimately detrimental to long-term host population viability if mutualisms are lost or decline in the future. Given the importance of mutualisms for biodiversity and ecosystem function^60,61^ as well as how mutualisms ameliorate stress for a diversity of organisms facing rapidly changing conditions^62–65^, this possibility is concerning. Therefore, we advocate for expeditious research investment by the scientific community into the interactive effects of changes in mean and variability of climate across a range of mutualisms that span the tree of life. These efforts may be vital for not only predicting how complex, real-world global change scenarios will affect the stability of these important stress-ameliorating interactions and the persistence of partnering organisms, but also for developing management strategies to protect the critical ecosystem services mutualisms underpin.

## Methods

### Study system

Endophytic fungi are ubiquitous, occurring in every major plant lineage^30^, with clavicipitaceous endophytes living as endosymbionts in the foliar tissue of an estimated 20–30% of the over 10,000 grass species (Poaceae)^66^. Occurrence estimates of systemic fungal endophytes of the genus *Epichloë* (Clavicipitaceae) in groups of cool-season grasses have ranged from 7.5%– 42.5%^31–33^. *Epichloë* endophytes are often mutualistic, conferring drought tolerance, resistance to herbivory and pathogens, and enhanced nutrient uptake to their hosts in exchange for photosynthetic carbon^23–25,67^. However, the costs and benefits of endophyte mutualism can vary through time and with environmental conditions^68^. This mutualism is facultative from the perspective of the plant and obligate from the perspective of the endophyte in that endophytes are not free-living and often depend on their hosts for transmission through the maternal lineage^30^.

*Bromus laevipes* (Shear) (Chinook brome) is a perennial C_3_ bunchgrass that occurs in small, patchy populations across the California floristic province^69^. *B. laevipes* commonly associates with vertically transmitted systemic fungal endophytes^34^. Previous research combining range-wide field surveys across 92 populations, species distribution modeling, field common garden experiments, and greenhouse experiments demonstrated that the presence of these endophytes was associated with the expansion of the geographic range of *B. laevipes* by thousands of square kilometers into drier habitats^13^. Furthermore, while endophytes confer drought tolerance, which is beneficial in drier environments, they likely also result in a net cost in wetter habitats, where carbon costs of supporting endophytes outweigh their drought amelioration benefits^13^.

### Population surveys

In 2009–2010, we surveyed endophyte prevalence in 92 natural populations of *B. laevipes* across northern and central California (19.81 ± 1.20 plants per population, mean ± SE). Sites were selected based on herbarium records and accessibility (*Consortium of California Herbaria, ucjeps*.*berkeley*.*edu/consortium*; see previous work^13^ for details on initial survey procedures). In 2022, we resurveyed ∼95% of the original 92 populations (15.94 ± 0.67 plants per population, mean ± SE, >1000 total; six sites not resurveyed due to inaccessibility; Figure 1), evaluating persistence of each population and quantifying endophyte prevalence (mutualism interaction frequency). Note *B. laevipes* is a short-lived perennial, with turnover of individuals expected between initial surveys and resurveys. To match previous survey methods for endophyte prevalence, each plant from a given population was evaluated for fungal hyphae by staining with aniline blue-lactic acid dye under a compound microscope (Figure S1), which yields similar results to immunoblot assay or PCR detection methods^70,71^ (see Supplementary Methods: *Seed staining for endophyte detection*). Population-level endophyte prevalence was then quantified as the percentage of individuals sampled from each host population in which endophyte was detected.

### Environmental data

To gain insight into the relationship between changing climates and both plant population persistence and mutualism interaction prevalence, we obtained data on two major environmental factors often shaped by climate change—aridity and fire history—for all surveyed grass population sites. For aridity/drought, downscaled climate data were obtained from the PRISM Climate Group for the period between the plant–endophyte surveys in 2009 and 2022. Then, potential evapotranspiration was calculated at each site using the *hargreaves* function, and the standardized precipitation evaporation index (SPEI) was calculated for each site using the *spei* function^72^. SPEI is a standardized drought index quantifying relative aridity of a site, with more negative values indicating more extreme drought and more positive values indicating wetter conditions^36^. We calculated three aspects of SPEI for each site over the period between surveys: mean SPEI (mean of the annual mean SPEIs), interannual variation in SPEI (standard deviation of annual mean SPEIs), and the rate of change in SPEI across those years (i.e., the regression coefficient between annual mean SPEI and time). For fire history, fire data were obtained from the California State Geoportal, and sites were scored dichotomously (1, 0) based on whether a fire had occurred during the period between the plant–endophyte surveys in 2009 and 2022. Data preparation and all statistical analyses were performed within R version 4.3.1 (R Core Team, 2023).

### Statistical analyses: Long-term population persistence and changes in mutualism prevalence in natural host populations

To determine if historical mutualism prevalence predicts plant population persistence and whether this is moderated by environmental factors, we first used global model selection to identify the best-supported multiple logistic regression model using the *glm* function. *B. laevipes* population persistence was the binary response variable. The candidate explanatory variables were historical endophyte prevalence (in 2009), mean SPEI, interannual variation in SPEI, rate of change in SPEI, and occurrence of fire between surveys, as well as the two-way interactions between historical endophyte prevalence and each environmental variable. Collinearities were checked using the *cor* function for pairs of continuous variables and by calculating an R^2^ value for pairs of continuous and categorical variables, with cutoffs of 0.7 and 0.49, respectively. We did not detect collinearities between candidate explanatory terms. Subsequent global model selection based on the corrected Akaike Information Criterion (AIC_c_) was conducted using the *dredge* function in R^73^ (Table S4). The best supported model had two explanatory variables, historical endophyte prevalence and fire occurrence. We further tested for an interaction effect in our best model (see Supplementary Methods: *Testing for interaction effects in the population persistence model*).

In order to determine which environmental factors promote or disrupt endophyte mutualism, we first evaluated whether host populations had shifted towards increased endophyte prevalence between surveys. Upon finding they did not (see Supplementary Discussion: *Host populations maintained intra-population variation in mutualism rather than progressing towards endophyte fixation*), we analyzed how endophyte prevalence shifted over time within populations with different environmental conditions and mutualism backgrounds. Specifically, we first constructed a multiple beta regression model using the *betareg* function^74^ with a response variable of current endophyte prevalence as a proportion (frequency in 2022) and candidate explanatory variables of historical endophyte prevalence, fire occurrence, and mean, variation, and rate of change in SPEI as well as the interactions between historical endophyte prevalence and each environmental variable. This was followed by global model selection (Table S6).

Residuals from all best models were checked for spatial autocorrelation using the *moran*.*test* function or 9999 permutations and the *moran*.*mc* function to conduct a permutation test for Moran’s I, depending on normality of residuals^75^. None of our statistical models exhibited significant spatial autocorrelation (P or pseudo-P > 0.05 for Moran’s I of all models).

### Demographic modeling to evaluate endophyte effects on predicted population persistence

In order to further assess and predict the effects of endophytes on plant population dynamics, we constructed a demographic model for *B. laevipes* parameterized with independent data from several multi-year, multi-site common garden experiments manipulating endophyte mutualism and measuring its effects on host plant recruitment, survival, and growth. We tested the predictions of our model against actual persistence of populations from our resurveys. Then, we evaluated our model across a set of abiotic conditions spanning the *B. laevipes* range and conducted life table response experiments to gain mechanistic understanding into how (i.e., through enhancing which vital rates) and under what conditions endophytes provide demographic benefits.

Because they predict population growth rates (λ), demographic model outputs are highly relevant to population persistence. In order to construct a population model, we first performed a set of common garden experiments tracking individual plants from 2010 to 2015. These experiments involved 1650 plants and 550 seeds, which were monitored for six years at five sites chosen to span a large part of the *B. laevipes* range^76^ (northern to central California; ∼420 km; Table S7), including a wide ecological and climatic gradient (e.g., ∼450–1750 mm average annual precipitation during experiment years, which includes 92% of the range of average annual precipitation experienced by this species during these years). To allow assessment of endophyte effects on host demography, fungal endophyte manipulation was achieved in two ways: first, by selecting plants and seeds from 11 populations with different endophyte mutualism statuses (i.e., naturally mutualistic and non-mutualistic populations) and second, through experimental fungicide application to allow assessment of endophyte demographic effects (i.e., experimental reduction of endophyte mutualism). (Please see Supplementary Methods: *Field common garden experiments for demographic model construction* for more details.) Using data from our experiments and from the PRISM database, we modeled vital rates (e.g., survival, growth, flower number) in relation to plant size and age, endophyte status, site and year-specific SPEI, and other factors (see Supplementary Methods: *Vital rate parametrizations*). We then used the vital rate models to parameterize a demographic model (see Supplementary Methods: *Population projection matrix model construction*). This model allowed us to predict growth rate for each host population site using its endophyte status and its yearly climate across the experimental years as inputs. For each population, we further predicted by how much the growth rate would increase if it was endophyte-associated compared to if it was endophyte-free, allowing us to relate endophyte population benefits to climatic factors.

Across the 86 natural populations we surveyed, we evaluated whether endophyte association was related to higher predicted population growth rates using 9999 permutations and the function *perm*.*t*.*test*^77^. We also performed fixed-effect life table response experiments to quantify class-dependent contribution of endophyte mutualism to the population growth rate^78^ (see Supplementary Methods: *Life table response experiment*). A life table response experiment calculates the contribution to population growth rate (i.e., difference in λ) of a fixed effect (here, endophyte mutualism) through all possible transitions between classes (Figure S12). We tested whether endophyte effects on population growth were greater via growth and survival or fecundity using a Wilcoxon signed-rank test (*wilcox*.*test* function). Finally, we evaluated how endophyte effects varied with climate conditions across our 86 field sites. For each site, we calculated the difference in λ between hypothetical endophyte-associated and endophyte-free populations, which represented the contribution of endophytes to population growth rates (i.e., the relative benefit of endophyte mutualism). We then ran a polynomial regression and a linear regression between this difference in modeled λ and the mean SPEI or variation in SPEI of each site across the experimental years, respectively, to determine whether endophyte mutualism effects on population growth rates varied by site climate.

## Data availability

All data and scripts will be uploaded to Zenodo upon acceptance of this manuscript.

## Author Contributions

VWL collected field survey data, performed data analysis and demographic modeling, and wrote the manuscript. JCF contributed to demographic model construction and manuscript revisions. ASD contributed to building and writing methods for the demographic model and manuscript revisions. SYS contributed to the conceptualization and design of the common garden experiments as well as the manuscript revisions and student supervision. CAS contributed to survey data collection, manuscript revisions, and feedback on modeling and statistical analyses. MEA led overall project conception and contributed to data collection for field surveys and common gardens, common garden experimental design and establishment, feedback on analyses, manuscript revisions, and student supervision.

## Acknowledgements

We thank the University of California Natural Reserve System, particularly the McLaughlin, Quail Ridge, Hastings, and Angelo Reserves, for providing protected natural habitats in which to conduct our experiments, the US Forest Service for their support of this project, and UC Reserve and Forest Service staff: J. Huhndorf, P. Aigner, C. Koehler, M. Power, J. Hunter, P. Steel, A. Spyres, R. Brennan, V. Boucher, J. Clary, L. Johnson, and M. Stromberg. We also thank the Afkhami and Searcy labs, D. Hernandez, A. O’Brien, W. Browne, and D. DeAngelis for helpful feedback on analyses and the manuscript. This research was supported by National Science Foundation support to MEA and CAS (NSF DEB-2030060 and NSF DEB-1922521).

