## Supplementary Information for "Climate variability disrupts mutualism-driven increases in population persistence"

### Supplementary Resources Contents

Supplementary Methods ........................................................ Pages 1 – 6
Supplementary Discussion ........................................................ Pages 7 – 9
Supplementary Figures (S1-S15) ........................................................ Pages 10 – 20
Supplementary Tables (S1-S13) ........................................................ Pages 21 – 32
Supplementary References ........................................................ Pages 33 – 34

### Supplementary Methods

*Seed staining for endophyte detection*

Each plant from a given population was evaluated for fungal hyphae by staining with aniline blue-lactic acid dye and examining leaf sheath tissue or seed tissue at 200x magnification under a compound microscope[1]. Leaf sheaths from three tillers (stored at 4ºC for up to 48 hours prior to staining) were examined per plant (if smaller than three tillers, all tillers were examined). For seeds, endophyte detection in at least two seeds was required to score the parent as endophyte-positive, while at least five seeds had to have no endophyte detected for the parent to be considered negative. If fewer than five seeds were available for a parent, one seed was sufficient to score the parent as endophyte-positive; this affected only five parental plants across our entire dataset (< 0.5% of all plants scored). No parents with at least five seeds had endophytes detected in only one seed.

*Testing for interaction effects in the population persistence model*

Global model selection identified two explanatory terms for our model of probability of population persistence, which were historical endophyte prevalence and fire occurrence. The interaction term was non-significant ($Z = 0.85$, $P = 0.40$) and was excluded through our model selection process. Because this was a generalized linear model (GLM), we further tested for an interaction effect by checking for a difference in the marginal effect of fire occurrence between 0% and 100% historical endophyte prevalence (following recommendations[2]) using the *second.diff.fitted* function[3]. We found that the effect of fire occurrence on predicted probability of population persistence did not depend on historical endophyte prevalence ($Z = 1.45$, $P = 0.15$).

Global model selection identified two significant explanatory terms for our model of current endophyte prevalence, which were historical endophyte prevalence and variability in aridity. Because this was a nonlinear model, we further tested for an interaction effect by checking for a difference in the marginal effect of variability in aridity between 0% and 100% historical endophyte prevalence (following recommendations[2]). We used the minimum and maximum variability in aridity values actually experienced by *both* historically highly mutualistic populations and historically non-mutualistic populations to perform this test. The *second.diff.fitted* function[3] used for our persistence GLM could not be applied to beta regression objects, so we manually tested for the significance of the interaction effect using 199 iterations of bootstrapping. We found that the effect of climate variability on predicted endophyte prevalence did depend on historical endophyte prevalence ($Z = -2.93$, $P = 0.0017$).

In addition to our beta regression, we considered several alternative frameworks to model endophyte prevalence using the same candidate explanatory variables and global model selection approach. Specifically, we fit the data using a multiple linear regression, polynomial regression, and binomial GLM. For the polynomial regression, the degree was determined using k-fold cross-validation and further verified using global model selection with candidate higher order terms for historical endophyte prevalence. Historical endophyte prevalence and variability in aridity were the only terms to be selected by the best model regardless of modeling approach, demonstrating their importance to current endophyte prevalence was robust. Furthermore, the interaction between historical endophyte prevalence and variability in aridity was always selected/significant. We ultimately chose to analyze and present the beta regression framework because of its ability to model proportional data without unevenly weighting *B. laevipes* populations.

*Field common garden experiments for demographic model construction*

To parameterize relevant vital rates to use to construct a demographic model, we conducted two sets of common garden experiments to monitor survival, growth, and reproduction of endophyte-associated and endophyte-free *B. laevipes* plants for six years. In January 2010, we established two sets of common gardens at five sites (10 gardens total) spanning a wide ecological and climatic gradient (e.g., ~450–1750 mm average annual precipitation during experiment years) and a large geographic range (northern to central California; ~420 km; Table S7)[4]. For the first set of common gardens, we germinated seeds sourced from 11 *B. laevipes* populations (Table S8) of varying levels of natural endophyte prevalence (five endophyte-associated and six endophyte-free) in the greenhouse, and seedlings from each of these 11 populations were transplanted into each site (330 seedlings per site; 1650 seedlings total). Endophyte-associated populations included three intermediate prevalence and two fixed (prevalence $\geq$ 90%) populations. The inclusion of intermediate frequency populations means some individuals from endophyte-associated populations are likely to be non-mutualistic, such that estimates of endophyte effects on population dynamics are likely conservative.

Prior to planting the first set of gardens in the field, we also experimentally treated half of the seeds from each of the five endophyte-associated populations with 2 g/L Benomyl fungicide following methods previously used to effectively manipulate endophyte infection in this[4,5] and other[6] grass species, thus allowing us to control for inherent or genotypic differences in demographic rates of plants from endophyte-associated and non-endophyte populations. Half of the seeds from the six endophyte-free populations received the same treatment to control for direct fungicide effects on host performance or on other fungi. Seeds were cold stratified at 4 °C for 2 weeks, then placed on a sunny laboratory bench for ~ 5 weeks to allow germination and initial growth. One-tiller seedlings from each treatment combination were transplanted into random gridded locations in the common gardens with 15 cm between individuals (a typical density for this plant species in nature). The seedlings were planted into an existing matrix of competitors under partial shade and partial sun at forest edges and experienced minimal disturbance. We collected data on survival, number of tillers produced, number of flowers, seeds produced, and damage per tiller from herbivory for all plants across six years. We defined dormancy in years 1–4 as those plants within a given year that appeared dead during sampling but were recorded as alive in subsequent years. Because of our inability to distinguish between dormancy and death during the final transition year of the experiment, we constructed our model only from the first four annual transitions.

We also conducted a second series of field germination common garden experiments using seeds from the same five naturally endophyte-associated populations and six naturally endophyte-free populations. At each of the five sites used in the common garden experiment, we established randomized planting arrays of seeds from all 11 populations (5 gardens × 11 populations × 10 seeds per population = 550 total seeds). Following previously used methods[5], we then tracked individual seed germination status for three years (2010–12). Because most germination occurred by the first year (76.91% of all seeds; 95.27% of seeds that germinated at any point), we treated the Year 1 data as the first year germination rate, which we differentiated from the seed bank that we define as viable seeds that have been in the ground for >1 year. We assumed that seeds that had not germinated after three years (19.27% of all seeds) were no longer viable. Because survival was modeled based on our transplant common garden experiments, predicted population growth rates could be inflated to some degree if survival of germinants from the germination experiments was lower than survival of small seedlings from the common garden transplant experiments, which had already germinated and achieved initial growth prior to planting in the field. To minimize inflated estimates of survival rates and subsequently population growth, we planted small (i.e., 1 tiller) seedlings at the initiation of the seedling common garden experiments.

*Vital rate parametrizations*

To construct our demographic model, we parameterized 12 vital rates using data from the common garden experiments, germination experiments, and the PRISM Climate Group database. These vital rates were survival probability, growth, flowering probability, flower number, seeds per tiller, first year germination probability, size of germinants, probability of entering dormancy, probability of leaving dormancy, size out of dormancy, probability of seed survival in seedbank, and probability of germination from seedbank (Table S9).

In order to account for differences in abiotic conditions between gardens, we included yearly site SPEI in our candidate vital rate models such that plants sampled in different combinations of gardens and years would receive unique SPEI values. Because SPEI is a standardized index, numerical SPEI values are not inherently meaningful, but rather reflect relative differences in the set of climatic measurements used to calculate them. In other words, equivalent SPEI values calculated from two different sets of climatic data could represent different climatic measurements. Thus, although five sites were involved in the common garden experiments, we calculated SPEI values across all 86 resurveyed sites to ensure that the full range of climatic measurements was included and all SPEI values were standardized relative to each other, which allowed us to later apply our model across all sites rather than just the five common garden sites. We also included measurements of herbivore damage in our vital rate functions to account for the potential influence of this important biotic factor.

We parameterized survival probability, flowering probability, probability of entering dormancy, and probability of leaving dormancy as binomial generalized linear mixed models (GLMMs), and growth, flower number, size of germinants, and size of dormancy as negative binomial GLMMs[7]. The vital rates modeling count data (e.g., number of flowers, or number of tillers for the growth model) were modeled using the negative binomial distribution to account for overdispersion. The variation (as standard deviation) for growth, size of germinants, and size out of dormancy was modeled using the generalized additive models for location, scale and shape (GAMLSS) framework[8]; for the growth model, standard deviation was explicitly modeled as a function of covariates. We modeled each vital rate and growth variance as functions of size, SPEI, herbivory damage, endophyte status, fungicide, and age as fixed effects and plant source population and common garden site as random effects. To account for variation in vital rates between populations, the population random effect was included to average across and account for genotype differences between populations. Furthermore, we used fungicide application to disentangle the effects of endophyte mutualism from potential population differences. In some systems, the effects of individual heterogeneity on population models can be small, especially once factors such as size and age are accounted for[9]. Thus, following previous demographic modeling research[10,11], we did not include individual as a random effect, although we tested its inclusion, which led to issues with model convergence during bootstrapping. When included, the individual random effect only improved the fit of one vital rate function (flowering probability; Table S10). Furthermore, while including the individual random effect consistently led to lower $\lambda$ predictions ($P < 0.0001$; permutation t-test; including individual random effect $1.37 \pm 0.027$ versus excluding individual random effect $1.44 \pm 0.026$; mean ± SE; Figure S5), the differences were relatively small (~5%) and did not qualitatively change the results or conclusions of any downstream analyses (e.g., comparing growth rates of endophyte-associated and endophyte-free populations). For each vital rate, we first constructed a global model and then reduced it through AIC-based backward stepwise model selection. In addition to the vital rate functions, we estimated three scalars using mean values: seed survival (probability that a seed has not germinated but is still viable), seed recruitment from the seed bank (i.e., germination in years 2 and 3), and the probability of leaving dormancy.

*Population projection matrix model construction*

Using the vital rate models we parameterized, we constructed a population projection matrix model to understand how endophyte mutualism affects population persistence through changes in plant demographic rates across the host range. We constructed a population projection matrix model over an integral projection model[12] in order to implement vital rate functions based on discrete probability distributions for count data.

Our model involved constructing a transition (growth and survival) subkernel P and a fecundity subkernel F from vital rate parameterizations. The transition subkernel P describes probabilities of an individual transitioning from one class (here, a given size and age) to another at a subsequent time point:

$$P(y_i,x_j) = s(x_j) \times g(y_i,x_j)$$

where $x_j$ is the class of an individual at time t, $y_i$ is the class of the individual at time t+1, $s(x_j)$ is the probability of survival for an individual of class $x_j$, and $g(y_i,x_j)$ is the probability that an individual of class $x_j$ will transition to class $y_i$ given that it survives.

The fecundity subkernel F describes the expected number of offspring for an individual of a given class. More specifically, it describes the probabilities of an individual of a given class x at time t producing offspring of another class y at time t+1, weighted by the predicted number of offspring:

$$F(y_i,x_j) = r_n(x_j) \times f_n(x_j) \times a_n(x_j) \times pE \times d(y_i)$$

where $r_n(x_j)$ is the class-specific probability of flowering, $f_n(x_j)$ is the class-specific number of flowers produced given that the individual has flowered, $a_n(x_j)$ is the class-specific number of seeds produced per flowered head, pE is the probability that a seed germinates and establishes, and $d(y_i)$ is the probability distribution of seedling size.

The transition and fecundity subkernels are then combined into a single matrix, the kernel K:

$$K(y,x) = P(y,x) + F(y,x)$$

Thus, the kernel essentially represents the transition probabilities over the range of classes of the organism of interest through both survival/growth and reproduction. In turn, population models can predict the population structure n (i.e., counts of individuals of different classes) at time t+1 by summing the product of the population structure at a previous time t and the kernel over the range of classes Ω, where x is the class at time t and y is the class at t+1:

$$n(y,t + 1) = \sum_{\Omega}[K(y,x)\ n(x,t)]\ dx$$

By varying the underlying factors of interest (e.g., endophyte mutualism, climate, etc.), vital rate functions (e.g., $r_n(x_j)$, $f_n(x_j)$) specific to each set of environmental conditions and/or population status result and different kernels can be constructed. This allows the population model to be applied to *B. laevipes* populations with different backgrounds of endophyte mutualism and predicts how they will perform in different environmental conditions.

Because age/year and fecundity were conflated (e.g., individuals by year four had substantially higher fecundity than individuals in year one, which were recently planted; Figure S6), we built a model that was both age- and size-structured (i.e., class consisted of a combination of age and size). The transition and fecundity subkernels were constructed for size classes of 1–17, where 17 represented the maximum observed tiller counts per individual in the dataset (15 tillers) times 1.1 and rounded up (following recommended approach[13]). The first two rows and columns of the transition and fecundity subkernels were designated for the seed bank and dormancy, respectively. These rows were filled using parameterized vital rate functions based on previous approaches[13]. The remaining elements (beginning with the third column and third row) represented individuals of size classes 1–17. 20 additional rows and columns were added to correct for potential eviction[12,14]. Four age classes were considered, one for each annual transition included in our model. In total, the transition and fecundity subkernels and the resulting kernel contained 39 x 39 x 4 matrix elements. However, our model predictions were robust regardless of whether or not eviction was accounted for (Figure S7).

While implementing our population model for each of our 86 sites based on available climate data and endophyte prevalence from our 2009 field surveys (which occurred the year prior to the initiation of our common garden experiments), we validated our model in two ways. First, we tested whether our model predicted higher population growth rates for populations observed to persist over the 13 year study period compared with populations that did not persist. This was accomplished using a permutation t-test implemented with function *perm.t.test* with 9999 permutations[15] (Figure S8). We further corroborated our model by testing whether population growth rates predicted for endophyte-free populations were equivalent to those predicted for endophyte-associated populations to which fungicide had been applied to remove the endophyte (see Figure S9). We ran a factorial ANOVA with site as a blocking term to test for differences between predicted growth rates for populations across the abiotic conditions spanning the *B. laevipes* range with different endophyte and fungicide statuses. Predicted growth rates for endophyte-free populations untreated with fungicide were significantly greater than ($P < 0.0001$) but were comparable to those of endophyte-associated populations treated with fungicide (endophyte-associated and fungicide $1.41 \pm 0.020$ versus endophyte-free and no fungicide $1.32 \pm 0.018$; mean ± SE). This demonstrated that fungicide treatment was partially effective in removing endophytes, which was in turn responsible for lower predicted $\lambda$ values. We also found that fungicide treatment lowered population growth rates in endophyte-associated populations ($P < 0.0001$; endophyte-associated and no fungicide $\lambda = 1.62 \pm 0.018$ versus endophyte-associated and fungicide $\lambda = 1.41 \pm 0.020$; mean ± SE). Because the endophyte-associated populations used in our common garden experiments to build vital rate functions were identical between the two fungicide treatments, this demonstrates that endophyte mutualism, and not the underlying host genotype, enhances host population growth.

*Life table response experiment*

To quantify class-dependent contributions of endophyte mutualism to the population growth rate through both class transitions and fecundity, we followed methods described previously[16]. More specifically, for each site of interest, we constructed transition or fecundity subkernels for both levels of endophyte status (i.e., endophyte-associated and endophyte-free). We constructed a third set of transition and fecundity subkernels averaged between endophyte statuses and constructed sensitivity matrices for both averaged subkernels based on previous approaches[13], which we then multiplied by matrices representing the difference between the endophyte-associated and endophyte-free subkernels[16].

### Supplementary Discussion

*Host populations maintained intra-population variation in mutualism rather than progressing towards endophyte fixation*

To evaluate whether or not populations trended towards fixation (i.e., 100% prevalence) of mutualism, we compared historical and current endophyte prevalence in populations with historically intermediate mutualism prevalence (n = 15 intermediate populations with 10-90% endophyte prevalence; cutoffs based on previous work[5]). We performed a linear regression of current endophyte prevalence (in 2022) on historical endophyte prevalence (in 2009). We determined if the intercept was significantly different from 0 or if a slope of 1 fell outside of the 95% CIs of the regression coefficient (calculated using the *confint* function), either or both of which would indicate current and historical endophyte prevalence are significantly different from a 1:1 ratio (i.e., mutualism prevalence in the populations changed significantly through time and thus mutualism moved towards either fixation or loss within populations).

As expected, historical endophyte prevalence and current endophyte prevalence were positively related ($R^2 = 0.54$, $F_{1,13} = 15.55$, P = 0.0017; Figure S13). However, ~25% of the mixed populations experienced complete endophyte loss, and no populations achieved 100% endophyte fixation. Furthermore, the regression line between current and historical endophyte prevalence did not differ significantly from the 1:1 line. The regression intercept (-25.66 ± 22.88; mean ± SE) did not differ significantly from 0 (T = -1.12, DF = 13, P = 0.28), and a slope of 1 fell within the 95% confidence interval of the regression coefficient (1.47 with 95% CI: [0.74,2.20]). Therefore, endophyte prevalence did not detectably change within historically mixed populations over the study period.

*The contribution of endophyte mutualism to population growth declines with increasing climatic variability*

To further explore how endophyte contribution to population growth interacted with climate, we created hypothetical locations spanning the climatic range and predicted population growth rates for each. These hypothetical locations each had different mean climates (i.e., SPEI values), corresponding to the range of observed mean climates in the 86 surveyed sites spanning the *B. laevipes* distribution during the experimental years. However, unlike our survey sites, SPEI for these hypothetical sites was held constant across all years, meaning climate variability for each site was 0. By then predicting λ for each hypothetical site, we were able to determine how population growth rates and endophyte mutualism's contribution to them was directly related to mean climate in the absence of temporal variability.

We found that while population growth rates of both endophyte-associated and endophyte-free populations increased with wetter conditions, suggesting drought is an important stressor for *B. laevipes*, endophyte-associated populations achieved optimal population growth rates under relatively drier conditions than endophyte-free populations (Figure S14). In turn, this difference in climatic optima meant the contribution of endophytes to population growth (i.e., their *relative* benefit; endophyte-associated λ minus endophyte-free λ) was greatest under intermediate to slightly dry climates (Figure S14). More climatically variable sites involve more extreme climatic years when conditions are much drier or wetter than average, and under such conditions our results indicate that the relative benefit of endophyte mutualism is reduced. Taken together, these results show the relative benefits of endophytes are greatest in average conditions and therefore decline with increasing climatic variability.

*Endophyte effects on host vital rates suggest possibilities for host–mutualist conflict*

Endophyte enhancement of host fecundity drove increased population growth rate predictions of host plants (Figure 3B). From our observations, *B. laevipes* plants tend to invest heavily in reproduction, so it was not surprising by itself that reproduction was the means through which endophytes benefitted host population dynamics. Surprisingly, however, endophytes did not benefit and in some cases even reduced host survival and growth, which could suggest subtle host–mutualist conflict. Vertically transmitted symbionts, such as these systemic fungal endophytes, are transmitted from host parent to offspring as opposed to between individuals within a generation[17]. In such symbioses, increased alignment of fitness interests between the vertically transmitted symbiont and their hosts resulting from coupled reproduction is expected to select for mutualistic interactions[18,19], and here, we found that endophytes indeed had a positive effect on host populations. However, even in co-evolved, vertically transmitted mutualisms, there are opportunities for meaningful conflict[20]. Interestingly, our life table response experiment revealed that endophyte demographic benefits largely occurred through enhanced host fecundity, whereas effects on host survival and growth were often negligible or even slightly negative. Potentially, fungal endophytes experience selection to benefit their hosts through fecundity even at a minor cost to survival and growth, because host reproduction—not survival and growth—is the means through which these endophytes are transmitted to new hosts. Many endophytes are transmitted uniparentally through the maternal lineage, and similar to how infected host plants exhibit bias in sex allocation towards the production of seeds over pollen[21], here, too, there is evidence that endophytes may shift host allocation in favor of investment into the mechanism they are transmitted through. This tendency for mutualists to shift host plant allocation in ways that increase their own fitness even at some cost to their host highlights subtle conflict between partners even in tightly coupled vertically transmitted mutualisms. However, this potential conflict between mutualist partners is masked by mutualist benefits to host fecundity, which dominate the net consequences of this interaction for host population dynamics in our field-parameterized models.

*Climate variability structures endophyte prevalence*

Endophyte prevalence within populations during our resurveys in 2022 declined with increasing climate variability (Figure 6). Interestingly, we found a similar pattern during our original surveys of *B. laevipes* populations in 2009. Specifically, endophyte prevalence in 2009 declined with increasing climate variability in an equivalent window of time prior to our original surveys, 1995–2008 ($R^2 = 0.27$, $F_{1,61} = 22.34$, $P < 0.0001$). Furthermore, we found that sites were more climatically variable in 2009–2022 than they had been between 1995–2008. Of the 63 populations that persisted to be resurveyed, only three (<5%) occupied sites that were less climatically variable in 2009–2022 than in 1995–2008, with the rest experiencing elevated climate variability (Figure S15). (Note that we calculated SPEI across the entire dataset, 1995–2022, before calculating SPEI SD to enable comparability between 1995–2008 and 2009–2022.) Therefore, we believe that climatic variability shaped endophyte prevalence in the past (i.e., leading up to our original 2009 surveys), has increased following the original surveys, and has likely since become even more important in structuring mutualism among these populations. This is reflected in our endophyte prevalence model (Figure 6)—after accounting for historical endophyte prevalence (during our original surveys), which was in part a product of historical climate variability pre-surveys, climate variability between surveys (2009–2022) significantly affected current endophyte prevalence.

### Supplementary Figures

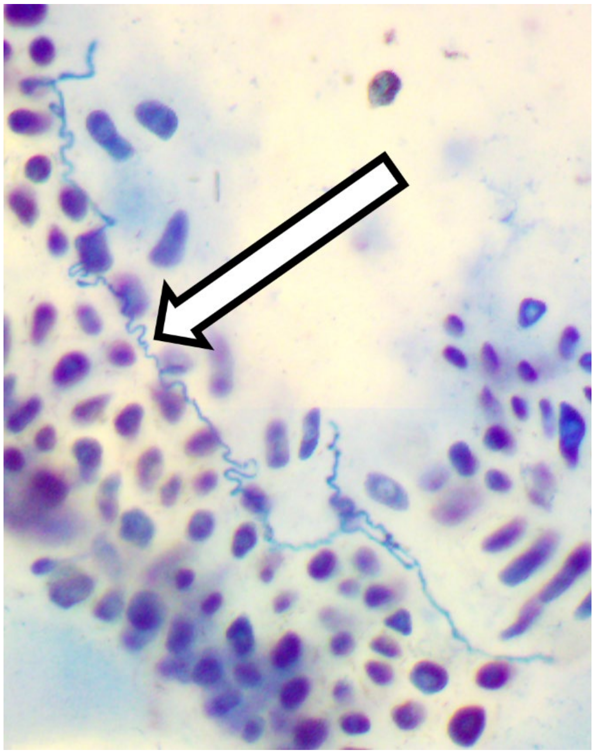

**Figure S1.** Lighter blue-stained fungal endophyte hyphae detected in the aleurone layer of a *B. laevipes* seed (200x magnification).

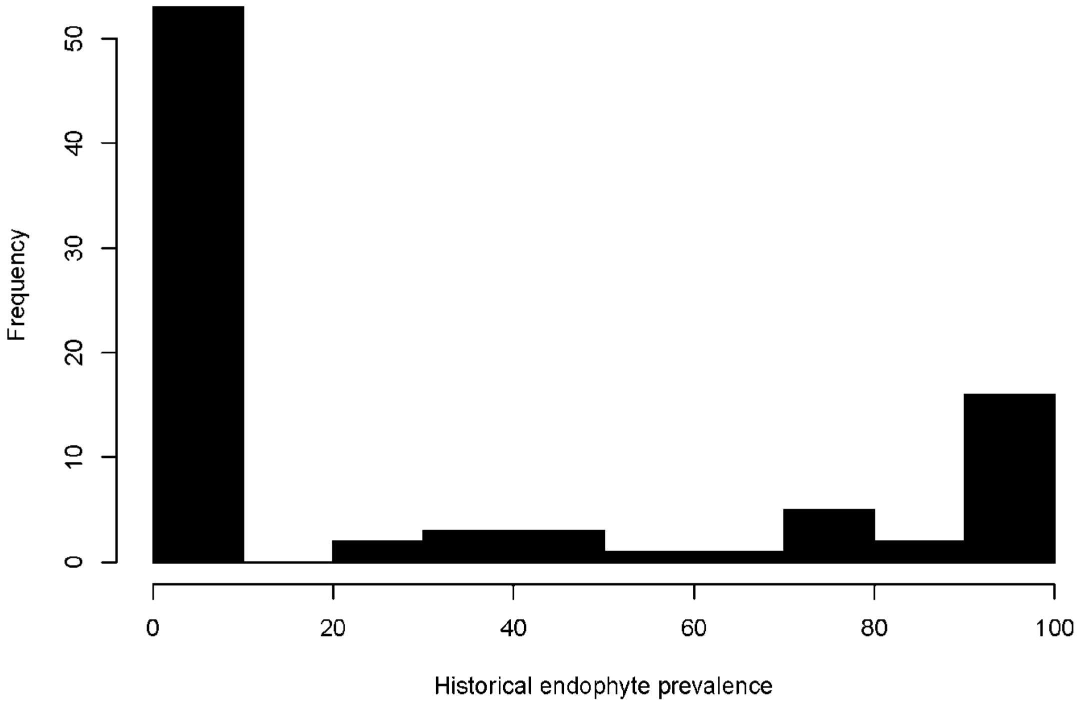


**Figure S2.** Histogram of historical endophyte prevalence (prevalence in 2009 during the original surveys ) in resurveyed *B. laevipes* populations.

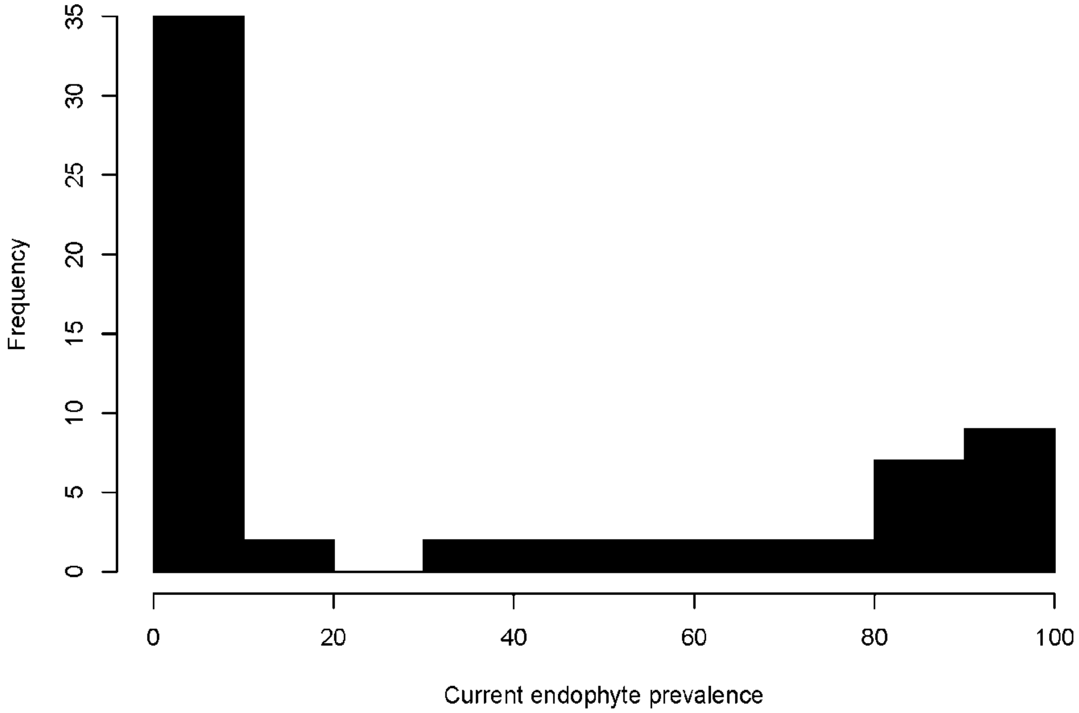


**Figure S3.** Histogram of current endophyte prevalence (in 2022 during the resureys) in resurveyed *B. laevipes* populations.

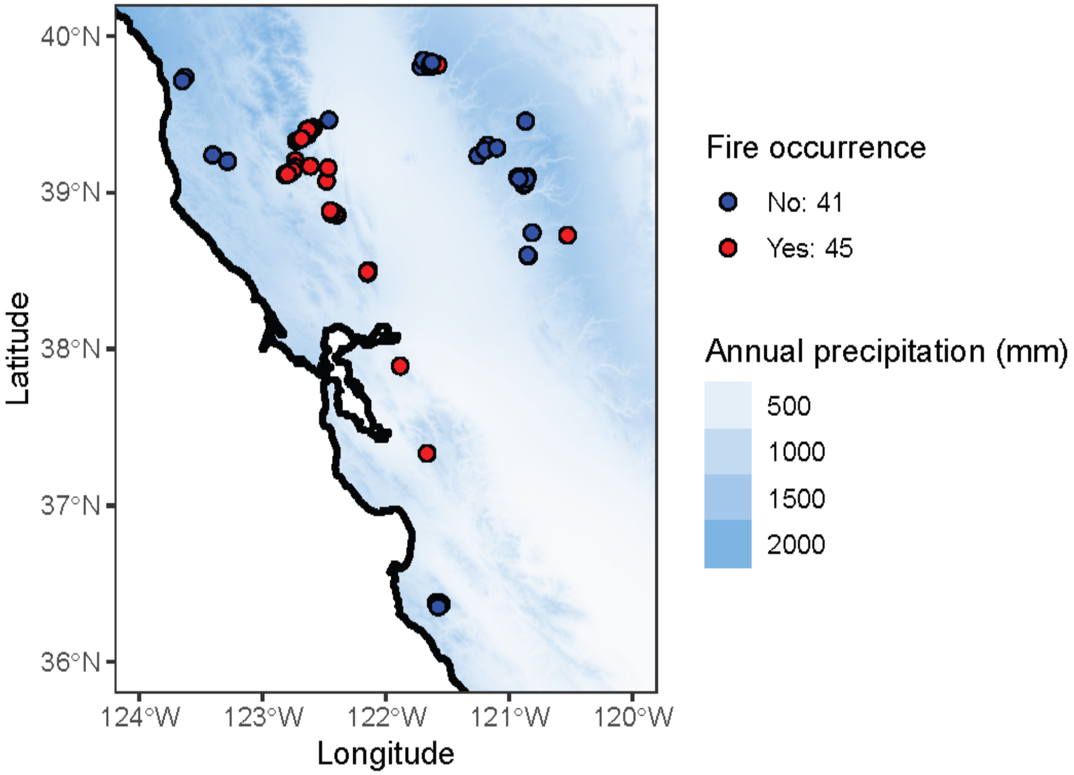

**Figure S4.** Many *B. laevipes* populations experienced fire between surveys and resurveys. Map colors represent annual precipitation (mm; data from WorldClim). Each point represents one population surveyed in 2009–10 and in 2022. Each point's color corresponds to whether or not it experienced fire between surveys and resurveys (data from the California State Geoportal), with 45 experiencing fire and 41 experiencing no fire.

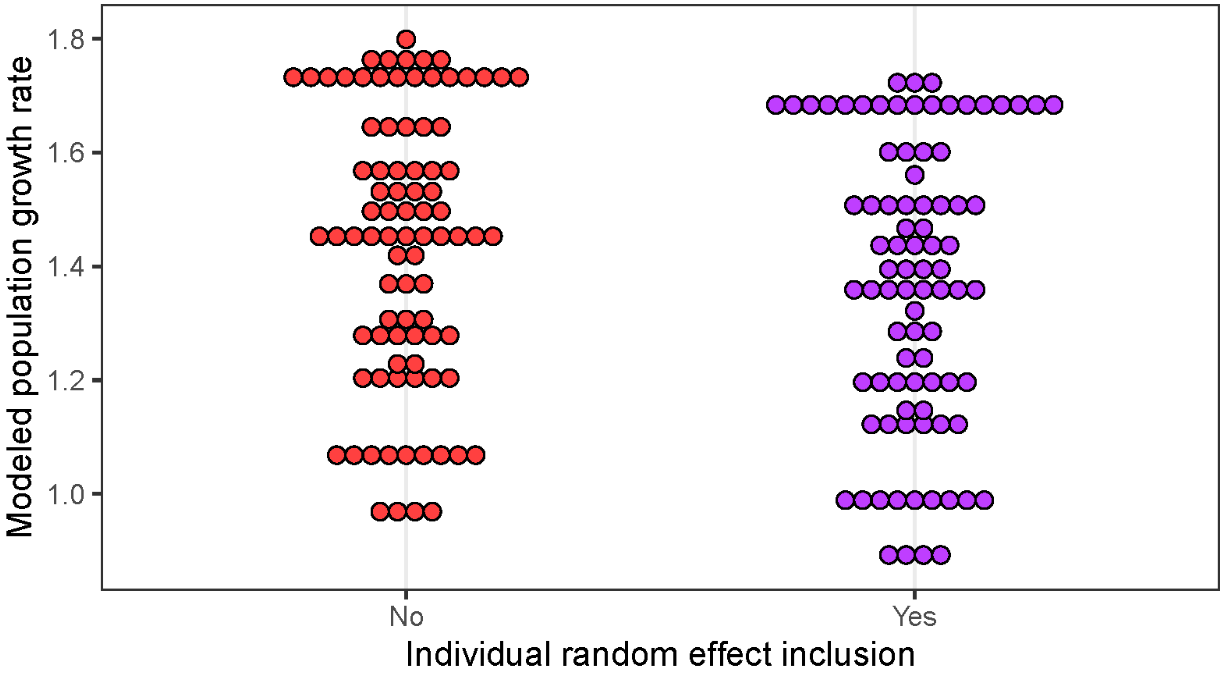


**Figure S5.** Modeled population growth rates for 86 resurveyed populations across the *B. laevipes* range. Including the individual host plant random effect in our vital rate models led to consistently lower $\lambda$ predictions for populations ($P < 0.0001$; including individual random effect $\lambda = 1.37 \pm 0.027$ versus excluding individual random effect $\lambda = 1.44 \pm 0.026$; mean ± SE). However, differences were small (~5%) and did not lead to any qualitatively different conclusions in downstream analyses.

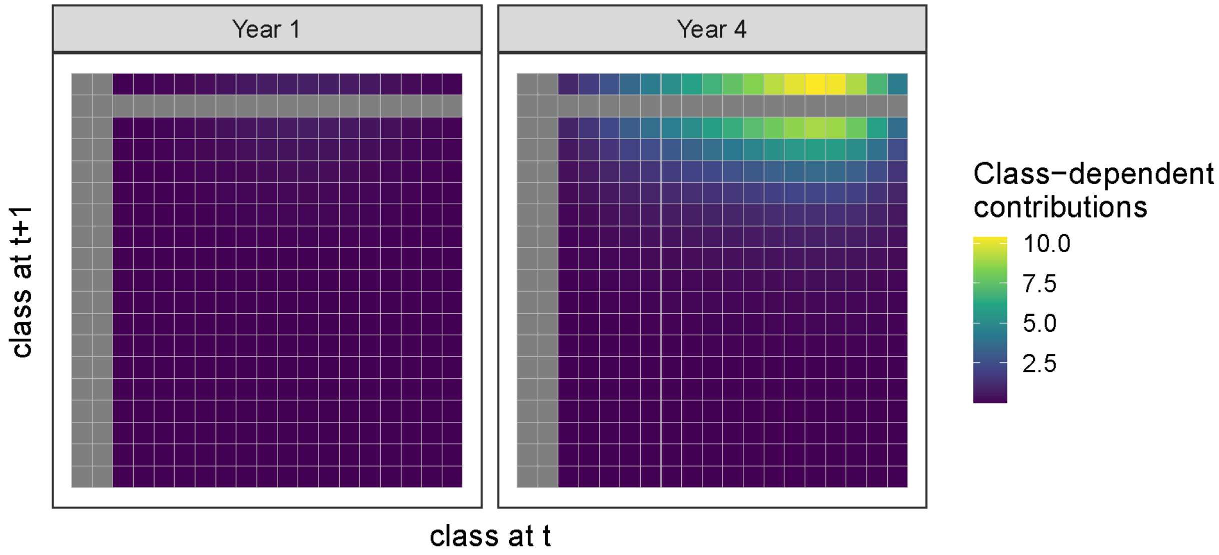


**Figure S6.** A visualization of a fecundity subkernel for a hypothetical population in a site with SPEI = 0 for all years of the experiment, corresponding to consistently “average” climatic conditions. Each element represents the contribution of an individual of a given class to another class in the next year through reproduction (i.e., the number of offspring of each class at time t + 1 produced, weighted by the probability of producing offspring, for a parent of a given class at time t). The first two classes (top- and left-most columns and rows) correspond to seeds and dormant individuals, respectively. The remainder of the rows and columns correspond to size classes 1–17 (i.e., individuals with 1–17 tillers). Additional size classes that were added to correct for eviction are not depicted here for visualization purposes. Gray cells represent 0 contribution to fecundity-based population growth. Because of these visually large differences in fecundity between ages, we constructed an age-structured population model.

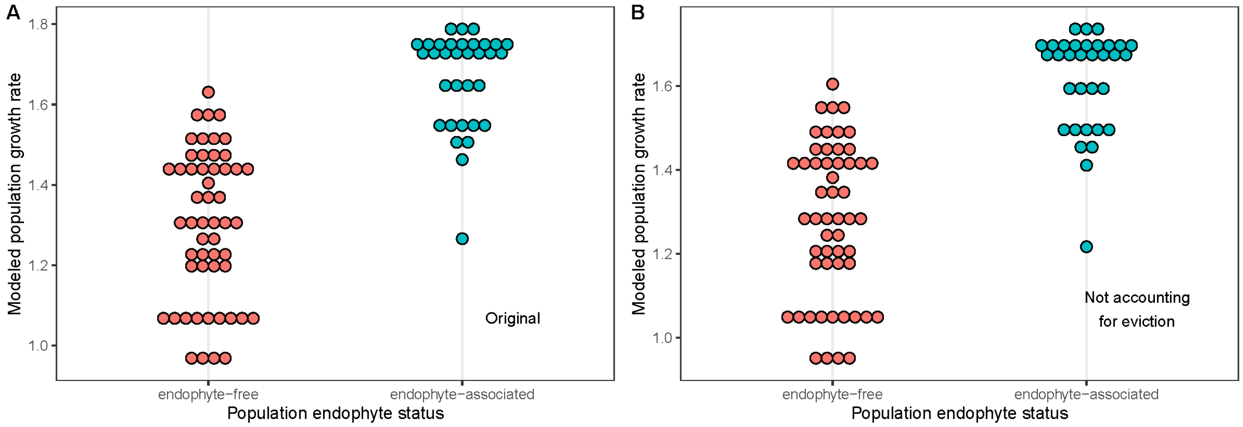


**Figure S7.** The transition and fecundity subkernels for our population projection matrix model had 20 additional rows and columns added to correct for potential eviction. However, model outputs were robust to whether potential eviction was considered **(A)** or not **(B)**, with population growth rates of endophyte-associated populations being significantly higher than endophyte-free populations. See main text for detailed results from models correcting for potential eviction. For models without correcting for potential eviction, population growth rates of endophyte-associated populations were also significantly greater than those of endophyte-free populations based on growth rates projected for each of the 86 surveyed populations (53 endophyte-free, 33 endophyte-associated; pseudo-P < 0.0001; $\lambda = 1.61 \pm 0.021$ for endophyte-associated populations vs $\lambda = 1.28 \pm 0.025$ for endophyte-free populations).

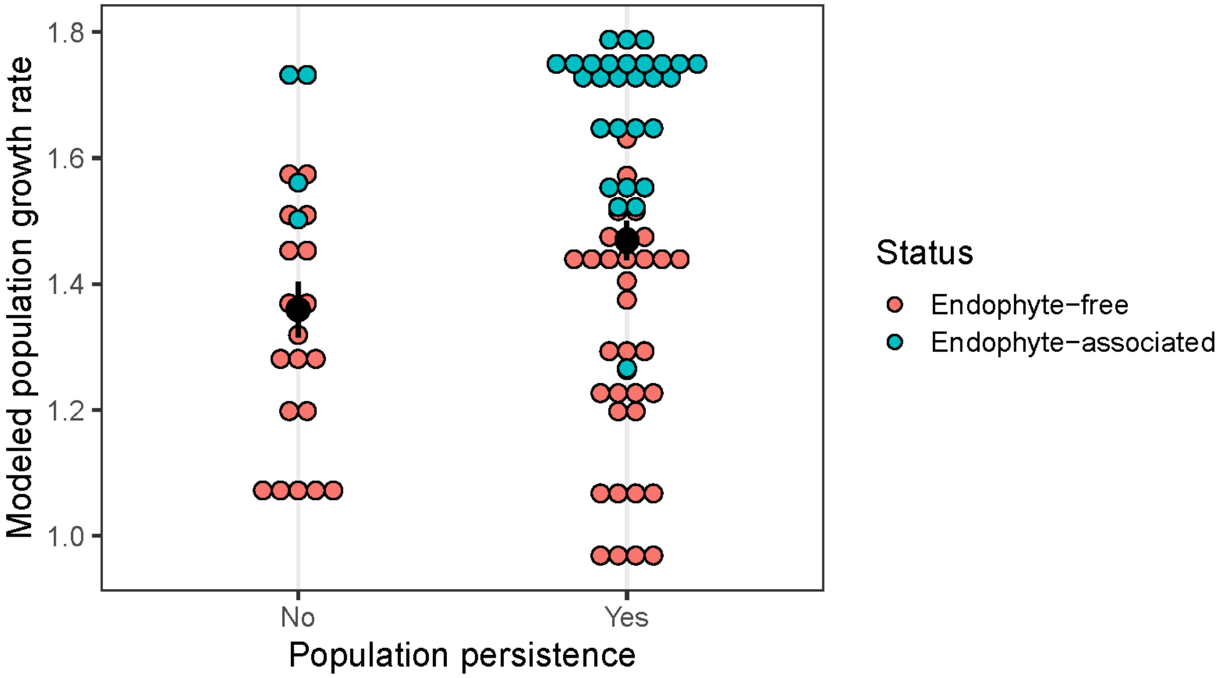


**Figure S8.** Modeled population growth rates for 86 resurveyed sites (63 persisted, 23 locally extinct as of resampling) across the *B. laevipes* range. Populations that persisted to resampling had marginally higher modeled population growth rates than populations that went locally extinct (pseudo-P = 0.062; 1.47 ± 0.031 for populations that persisted vs 1.36 ± 0.044 for locally extinct populations, mean ± SE). Black circles and lines represent mean and standard error values of predicted λ for each group. Point color represents endophyte status of each population; note that populations that persisted were more commonly endophyte-associated and tended to have higher predicted λ values (also see Figure 3A).

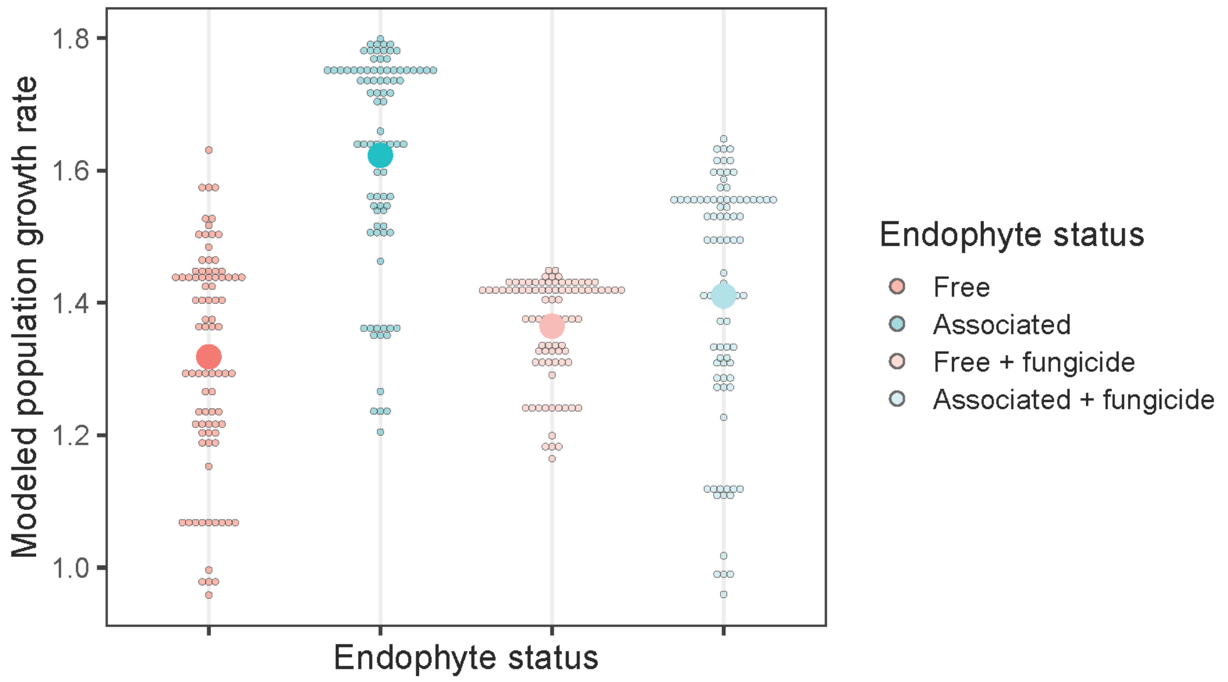


**Figure S9.** Modeled population growth rates for 86 resurveyed sites across the *B. laevipes* range including fungicide treatment as a potential term, demonstrating fungicide was largely successful in reducing endophyte-association effects on modeled population growth unrelated to genotype differences between endophyte-associated and endophyte-free *B. laevipes* populations. The large points represent the mean of each group's predicted population growth rates. Note that fungicide was not specifically targeted towards endophytic fungi and could have removed other symbiotic fungi as well, allowing for potential biological effects aside from purely removal of endophytes. Thus, we included an endophyte-free + fungicide treatment to control for non-target effects of fungicide.

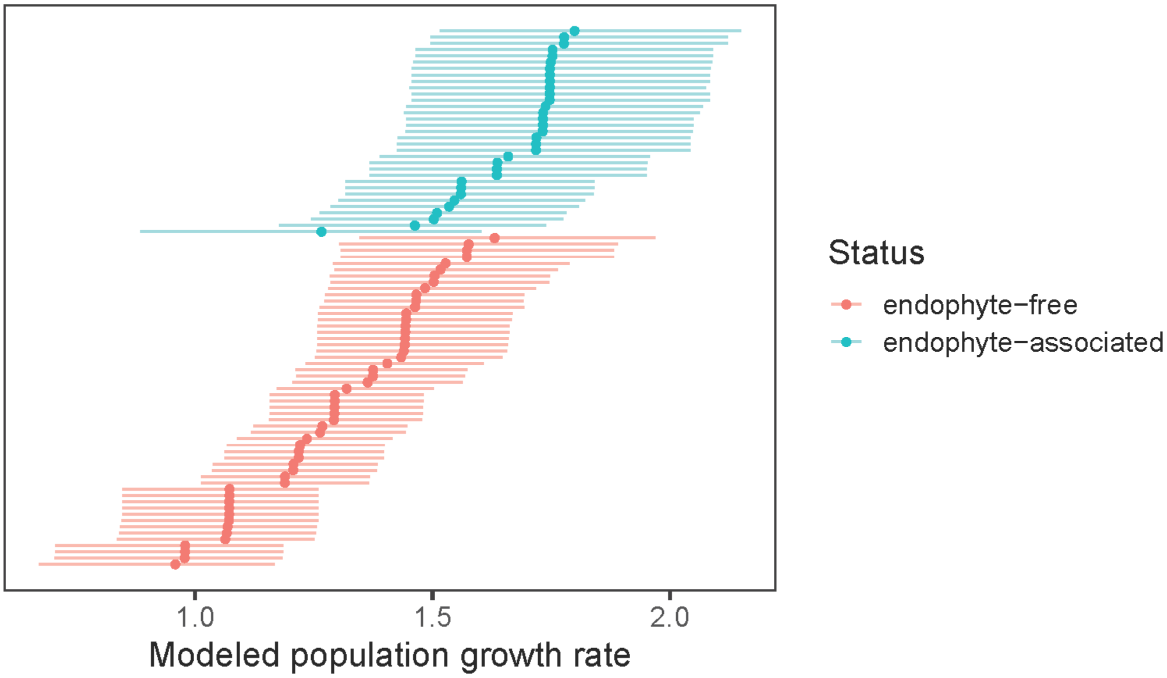


**Figure S10.** Modeled population growth rates for 86 resurveyed sites (53 endophyte-free, 33 endophyte-associated) across the *B. laevipes* range. Error bars represent bootstrapped 95% confidence intervals. Endophyte-associated populations had 28.37% higher predicted population growth rates on average than endophyte-free populations (pseudo-P < 0.0001; $\lambda = 1.67 \pm 0.021$ for endophyte-associated populations vs $\lambda = 1.30 \pm 0.025$ for endophyte-free populations; also see Figure 3A).

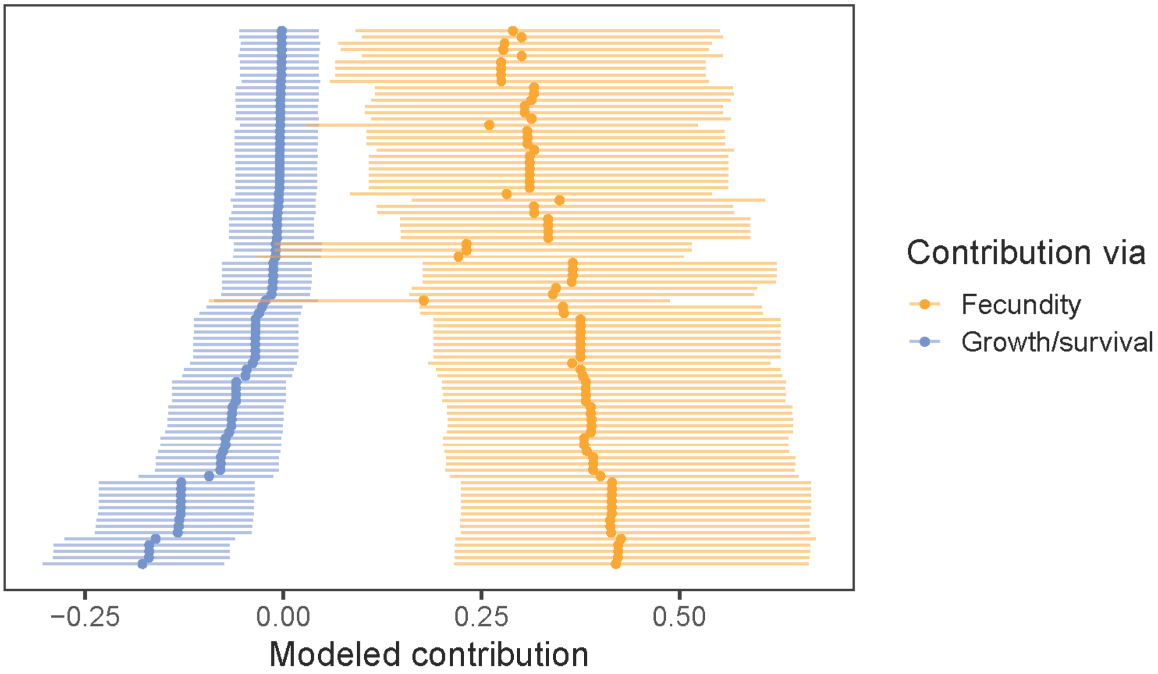


**Figure S11.** Modeled endophyte contributions to population growth rates for the 86 resurveyed sites across the *B. laevipes* range through vital rates related to host fecundity and survival/growth. Endophyte contributions to population growth rates were greater through fecundity than through survival/growth (Wilcoxon signed-rank test: V = 3741, Z = 8.05, P < 0.0001; mean ± SE contribution: -0.045 ± 0.0055 through survival/growth, 0.35 ± 0.0058 through fecundity; mean ± SE contribution difference: 0.39 ± 0.011; also see Figure 3B). Error bars represent bootstrapped 95% confidence intervals.

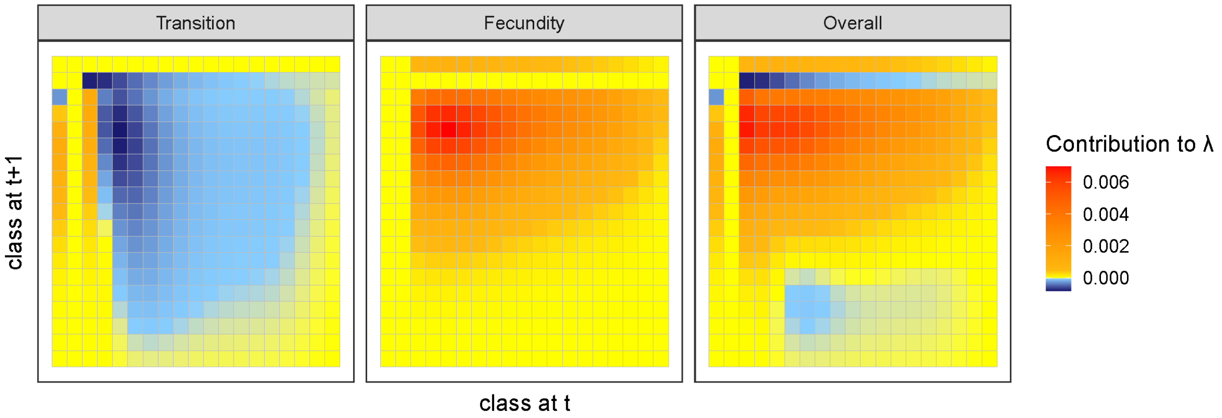


**Figure S12.** An example of the output of a life table response experiment for a population in a site that experiences stable and average aridity (i.e., occurs in a site with SPEI = 0 for all years of the experiment, corresponding to consistently “average” climatic conditions). Here, class-dependent transitions have been summed across all ages to create a two-dimensional matrix for visualization. Each element represents the contribution to population growth rate of endophyte mutualism through each size class’s vital rates (either 1. transition—the combined survival and growth subkernel, 2. fecundity, or 3. overall contribution), summed over all age classes in our model. The first two classes (top- and left-most columns and rows) in each heatmap correspond to seeds and dormant individuals, respectively. The remainder of the rows and columns correspond to size classes 1–17 (i.e., individuals with 1–17 tillers). Additional size classes that were added to correct for eviction are not depicted here for visualization purposes. In this example, most of the contribution of endophyte mutualism to population growth rate is through fecundity, specifically the production of relatively small offspring.

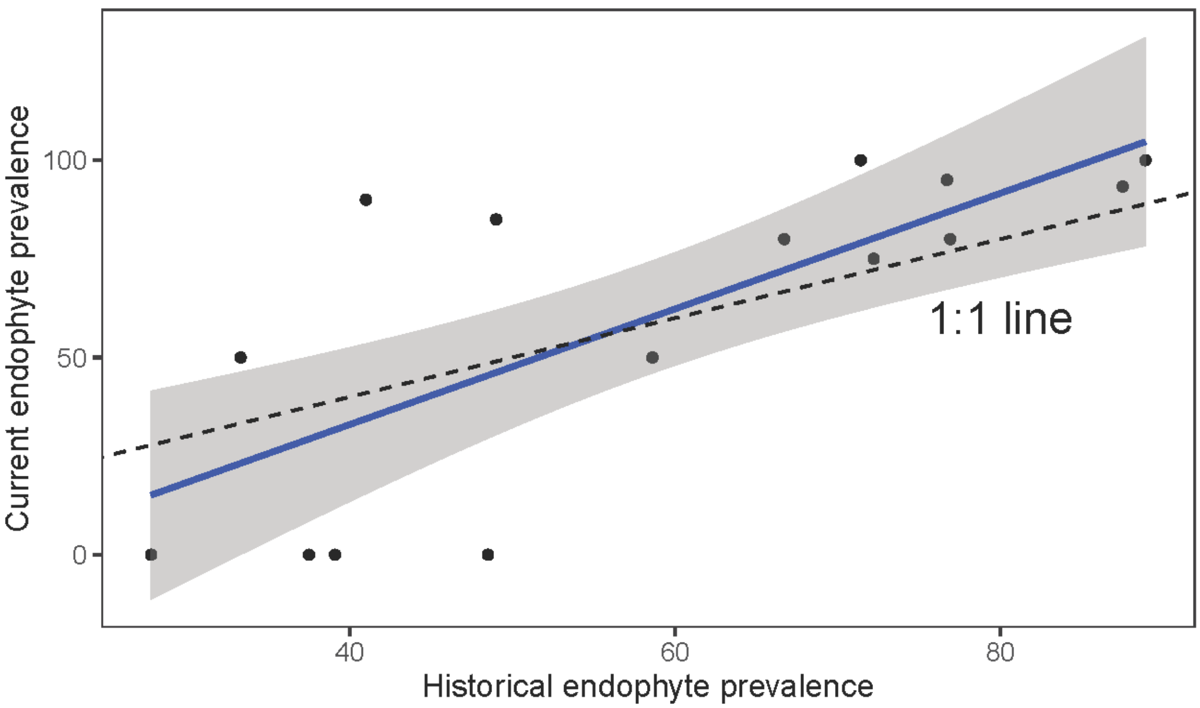


**Figure S13.** Current endophyte prevalence was positively correlated with historical endophyte prevalence for *B. laevipes* populations with historically intermediate endophyte prevalence ($R^2$ = 0.54, $F_{1,13}$ = 15.55, P = 0.0017). Here, current endophyte prevalence refers to endophyte prevalence in a given surveyed field population in 2022 and historical endophyte prevalence refers to its prevalence in 2009. Each point represents one *B. laevipes* population, the solid blue line is the linear regression, and the shaded areas represent the standard error. The black dashed line depicts the 1:1 line between current and historical endophyte prevalence. Contrary to expectations, because the regression coefficient (1.47 with 95% CI: [0.74,2.20]) was not significantly different from 1 and the intercept (-25.66 ± 22.88; mean ± SE) was not significantly different from 0, endophyte prevalence did not increase significantly towards fixation in populations with historically intermediate endophyte prevalence over the study period.

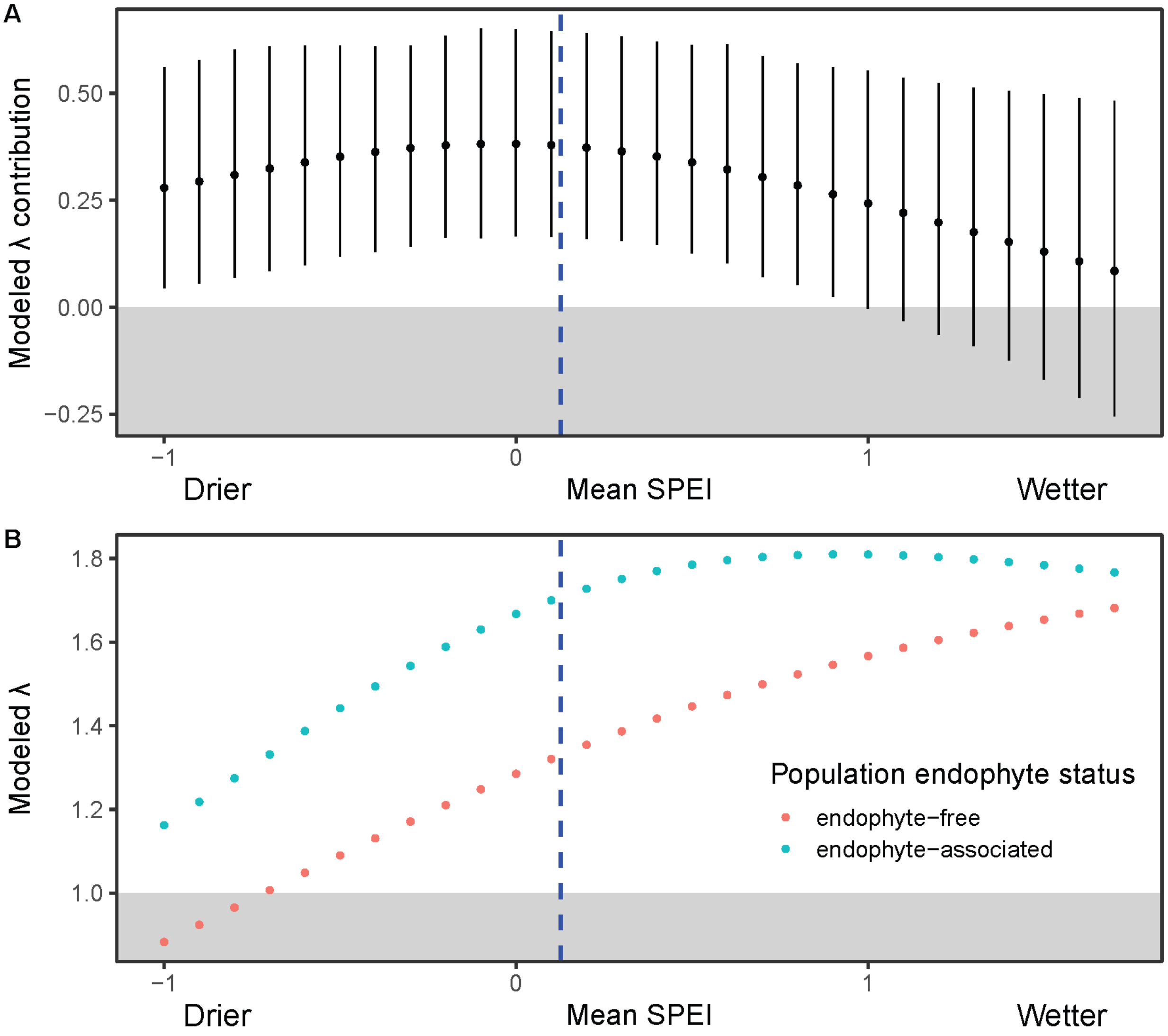


**Figure S14. (A)** Endophyte contributions to modeled population growth rates (i.e., the difference in modeled λ between endophyte-associated and endophyte-free populations) for hypothetical sites with different mean climates and no year-to-year variability in climate. Endophyte contributions are greatest under roughly average climates. Error bars represent bootstrapped 95% confidence intervals. **(B)** Modeled population growth rates for the same hypothetical populations. As sites became wetter on average during the four experimental years, predicted λ values increased for both endophyte-associated and endophyte-free populations. However, endophyte-associated populations achieve optimal population growth rates under drier on average conditions than endophyte-free populations. The dashed blue line represents the mean SPEI for all 86 surveyed sites of *B. laevipes* populations over the four experimental years; note that the mean is not centered at zero because the climate values from the four common garden experimental years represent only a subset of the climate data used to calculate SPEI values.

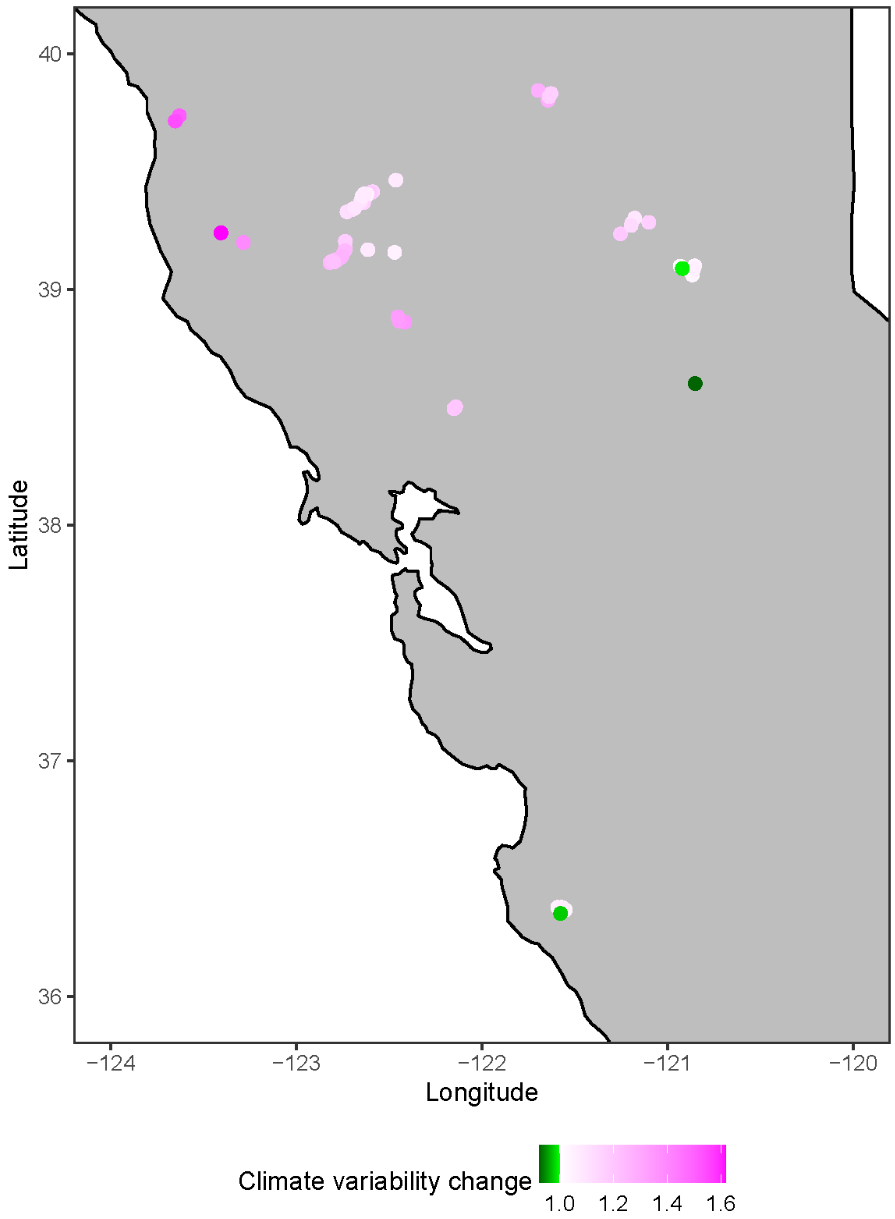


**Figure S15.** The vast majority of *B. laevipes* populations that persisted to be resurveyed in 2022 experienced increased climate variability in 2009–2022 (i.e., between surveys and resurveys) than they did prior to our original surveys in 2009. Each point represents one population surveyed in 2009–10 that persisted to resurveying in 2022. Each point's color corresponds to the ratio of its interannual variability in aridity between surveys (2009–2022) to its variability in an equivalent time window prior to the original surveys (1995–2008); note the color scale. Only three populations (< 5%; these populations are shown in green) experienced decreased climate variability between surveys compared to prior to surveys; most populations have experienced increases in climate variability.

### Supplementary Tables

**Table S1.** Locality information and endophyte prevalence for *Bromus laevipes* populations surveyed in 2009–10 and in 2022.

| Latitude | Longitude | Region | Historical endophyte prevalence (2009) | Current endophyte prevalence (2022) |
|---|---|---|---|---|
| 39.73073 | -123.641 | Coast | 0 | Population gone |
| 37.891 | -121.885 | Coast Range | 0 | Population gone |
| 39.40057 | -122.613 | Coast Range | 0 | Population gone |
| 39.4053 | -122.636 | Coast Range | 0 | Population gone |
| 39.07348 | -122.482 | Coast Range | 0 | Population gone |
| 37.33286 | -121.667 | Coast Range | 0 | Population gone |
| 38.85668 | -122.397 | Coast Range | 0 | Population gone |
| 38.48491 | -122.148 | Coast Range | 0 | Population gone |
| 38.48905 | -122.152 | Coast Range | 0 | Population gone |
| 38.4858 | -122.146 | Coast Range | 0 | Population gone |
| 38.48835 | -122.147 | Coast Range | 0 | Population gone |
| 38.4836 | -122.149 | Coast Range | 0 | Population gone |
| 39.04962 | -120.881 | Sierra Foothills | 0 | Population gone |
| 39.45577 | -120.868 | Sierra Nevada | 0 | Population gone |
| 39.80492 | -121.716 | Sierra Nevada | 0 | Population gone |
| 38.59353 | -120.845 | Sierra Nevada | 0 | Population gone |
| 39.81612 | -121.578 | Sierra Nevada | 0 | Population gone |
| 38.74338 | -120.815 | Sierra Nevada | 0 | Population gone |
| 38.72712 | -120.528 | Sierra Nevada | 0 | Population gone |
| 39.09867 | -120.924 | Sierra Nevada | 25 | Population gone |
| 38.86988 | -122.447 | Coast Range | 74.3 | Population gone |
| 38.87428 | -122.424 | Coast Range | 100 | Population gone |
| 39.08841 | -120.921 | Sierra Nevada | 100 | Population gone |
| 39.737 | -123.63 | Coast | 0 | 0 |
| 39.71472 | -123.652 | Coast | 0 | 14.29 |
| 39.405 | -122.636 | Coast Range | 0 | 0 |
| 39.41457 | -122.589 | Coast Range | 0 | 0 |
| 39.41305 | -122.597 | Coast Range | 0 | 0 |
| 39.39957 | -122.617 | Coast Range | 0 | 0 |
| 39.39321 | -122.63 | Coast Range | 0 | 0 |
| 39.40858 | -122.604 | Coast Range | 0 | 0 |
| 39.36814 | -122.638 | Coast Range | 0 | 0 |
| 39.36922 | -122.656 | Coast Range | 0 | 0 |
| 39.32948 | -122.727 | Coast Range | 0 | 0 |
| 39.1738 | -122.735 | Coast Range | 0 | 0 |
| 39.127 | -122.78 | Coast Range | 0 | 0 |

| | | | | |
|---|---|---|---|---|
| 38.8611 | -122.416 | Coast Range | 0 | 0 |
| 38.86595 | -122.446 | Coast Range | 0 | 0 |
| 38.50188 | -122.142 | Coast Range | 0 | 0 |
| 38.49371 | -122.151 | Coast Range | 0 | 0 |
| 39.06163 | -120.867 | Sierra Foothills | 0 | 0 |
| 39.84425 | -121.696 | Sierra Nevada | 0 | 0 |
| 39.80342 | -121.644 | Sierra Nevada | 0 | 0 |
| 39.80974 | -121.643 | Sierra Nevada | 0 | 0 |
| 39.09822 | -120.932 | Sierra Nevada | 0 | 0 |
| 39.09988 | -120.854 | Sierra Nevada | 0 | 0 |
| 39.81791 | -121.638 | Sierra Nevada | 0 | 0 |
| 39.8315 | -121.628 | Sierra Nevada | 0 | 0 |
| 39.28296 | -121.192 | Sierra Nevada | 0 | 0 |
| 39.30257 | -121.177 | Sierra Nevada | 0 | 0 |
| 39.23561 | -121.254 | Sierra Nevada | 0 | 0 |
| 39.2712 | -121.196 | Sierra Nevada | 0 | 0 |
| 39.08743 | -120.859 | Sierra Foothills | 0 | 14.29 |
| 39.14593 | -122.752 | Coast Range | 0 | 30 |
| 39.15739 | -122.47 | Coast Range | 0 | 40 |
| 39.28477 | -121.101 | Sierra Nevada | 0 | 85.71 |
| 38.60054 | -120.851 | Sierra Nevada | 0 | 0 |
| 39.34433 | -122.688 | Coast Range | 27.8 | 0 |
| 39.13443 | -122.764 | Coast Range | 33.3 | 50 |
| 38.8743 | -122.445 | Coast Range | 37.5 | 0 |
| 39.08865 | -120.92 | Sierra Nevada | 39.1 | 0 |
| 39.40397 | -122.619 | Coast Range | 41 | 90 |
| 39.46385 | -122.464 | Coast Range | 48.5 | 0 |
| 39.38088 | -122.651 | Coast Range | 49 | 85 |
| 39.39677 | -122.645 | Coast Range | 58.6 | 50 |
| 39.40227 | -122.631 | Coast Range | 66.7 | 80 |
| 36.36667 | -121.55 | Coast | 71.4 | 100 |
| 39.20421 | -122.736 | Coast Range | 72.2 | 75 |
| 39.33877 | -122.7 | Coast Range | 76.7 | 95 |
| 39.19951 | -123.285 | Coast | 76.9 | 80 |
| 39.12433 | -122.794 | Coast Range | 87.5 | 93.33 |
| 36.37754 | -121.592 | Coast | 88.9 | 100 |
| 36.37754 | -121.57 | Coast | 90 | 100 |
| 39.1196 | -122.791 | Coast Range | 94.7 | 85 |
| 38.88293 | -122.453 | Coast Range | 100 | 0 |
| 39.11361 | -122.82 | Coast Range | 100 | 5 |
| 39.12171 | -122.791 | Coast Range | 100 | 35 |
| 36.3711 | -121.565 | Coast | 100 | 45 |
| 39.24014 | -123.406 | Coast | 100 | 66.67 |

| | | | | |
|---|---|---|---|---|
| 39.16421 | -122.739 | Coast Range | 100 | 64.71 |
| 39.13981 | -122.757 | Coast Range | 100 | 75 |
| 39.1191 | -122.809 | Coast Range | 100 | 85 |
| 39.11663 | -122.799 | Coast Range | 100 | 95 |
| 36.35154 | -121.576 | Coast | 100 | 100 |
| 39.34477 | -122.686 | Coast Range | 100 | 80 |
| 39.1685 | -122.615 | Coast Range | 100 | 100 |

NOTE: All localities were unique populations separated by intervening habitat unsuitable for the host plant species.

**Table S2.** Number of *B. laevipes* populations with each historical endophyte status in 2009 divided by whether or not they persisted to resurveying in 2022 (fixed, prevalence $\geq$ 90%; intermediate, 10% < prevalence < 90%; non-mutualistic, prevalence $\leq$ 10%; cutoffs based on previous work[5]; also see Figure 1).

| | Persistence | Extinction |
|---|---|---|
| Fixed | 14 | 2 |
| Intermediate | 15 | 2 |
| Non-mutualistic | 34 | 19 |

**Table S3.** Statistical results for population persistence modeled as a logistic regression of historical endophyte prevalence and fire occurrence between surveys. This was determined to be the best-supported model through global model selection.

| | $\chi^2$ | DF | P |
|---|---|---|---|
| Historical endophyte prevalence | 6.73 | 1 | 0.0095 |
| Fire occurrence | 2.91 | 1 | 0.088 |

**Table S4.** Comparison of candidate models for population persistence. Each row represents a candidate model, ordered by $AIC_c$. The last three columns describe properties of each candidate model, with the remaining columns each representing an explanatory term in the model. NA values indicate the explanatory term was not present in the candidate model. Inclusion of a term in the model is indicated by + (for categorical variables) and numerical (for continuous variables, where the number is the coefficient estimate) values.

| Fire occurrence | Mean SPEI | Coefficient of regression of SPEI on time | Historical endophyte prevalence | SPEI SD | Endophyte :fire | Endophyte :mean SPEI | Endophyte :SPEI coefficient of regression | Endophyte :SPEI SD | DF | Log-likelihood | AICc |
|---|---|---|---|---|---|---|---|---|---|---|---|
| + | NA | NA | 1.69 | NA | NA | NA | NA | NA | 3.00 | -45.52 | 97.33 |
| + | NA | 0.51 | -4.00 | NA | + | NA | -5.00 | NA | 6.00 | -42.41 | 97.88 |
| NA | NA | NA | 1.56 | NA | NA | NA | NA | NA | 2.00 | -46.97 | 98.09 |
| + | NA | 0.41 | -1.05 | NA | NA | NA | -3.52 | NA | 5.00 | -43.83 | 98.41 |
| NA | NA | 0.12 | -1.17 | NA | NA | NA | -3.64 | NA | 4.00 | -44.98 | 98.46 |
| + | NA | 0.84 | 9.92 | 0.90 | + | NA | -5.50 | -15.39 | 8.00 | -40.36 | 98.59 |
| + | NA | NA | 0.91 | NA | + | NA | NA | NA | 4.00 | -45.17 | 98.84 |

| | | | | | | | | | | | |
|---|---|---|---|---|---|---|---|---|---|---|---|
| NA | NA | NA | 16.53 | 0.67 | NA | NA | NA | -14.41 | 4.00 | -45.18 | 98.86 |
| + | NA | NA | 16.09 | 0.58 | NA | NA | NA | -13.83 | 5.00 | -44.13 | 99.01 |
| + | NA | NA | 1.71 | 0.14 | NA | NA | NA | NA | 4.00 | -45.49 | 99.47 |
| + | -0.09 | NA | 1.70 | NA | NA | NA | NA | NA | 4.00 | -45.50 | 99.50 |
| + | NA | 0.08 | 1.66 | NA | NA | NA | NA | NA | 4.00 | -45.51 | 99.51 |
| + | NA | 0.69 | -1.69 | 0.68 | NA | NA | -4.36 | NA | 6.00 | -43.35 | 99.76 |
| NA | NA | 0.37 | -1.74 | 0.58 | NA | NA | -4.42 | NA | 5.00 | -44.60 | 99.95 |
| + | NA | 0.63 | -4.01 | 0.35 | + | NA | -5.28 | NA | 7.00 | -42.28 | 100.00 |
| + | NA | NA | 15.09 | 0.55 | + | NA | NA | -14.00 | 6.00 | -43.49 | 100.04 |
| NA | NA | -0.26 | 1.67 | NA | NA | NA | NA | NA | 3.00 | -46.88 | 100.05 |
| + | -0.19 | 0.52 | -4.01 | NA | + | NA | -5.03 | NA | 7.00 | -42.35 | 100.14 |
| NA | NA | NA | 1.60 | 0.16 | NA | NA | NA | NA | 3.00 | -46.94 | 100.17 |
| NA | 0.03 | NA | 1.56 | NA | NA | NA | NA | NA | 3.00 | -46.97 | 100.24 |
| + | -0.57 | 1.02 | 11.54 | 1.16 | + | NA | -5.73 | -17.23 | 9.00 | -39.95 | 100.26 |
| + | -0.16 | 0.42 | -1.07 | NA | NA | NA | -3.57 | NA | 6.00 | -43.79 | 100.64 |
| + | -0.27 | NA | 1.64 | NA | NA | 0.91 | NA | NA | 5.00 | -44.96 | 100.66 |
| NA | -0.02 | 0.12 | -1.18 | NA | NA | NA | -3.64 | NA | 5.00 | -44.98 | 100.72 |
| + | NA | 0.40 | 17.38 | 0.73 | NA | NA | NA | -15.20 | 6.00 | -43.95 | 100.97 |
| NA | NA | 0.54 | 8.21 | 0.93 | NA | NA | -3.08 | -8.97 | 6.00 | -43.98 | 101.01 |
| + | -0.13 | NA | 0.90 | NA | + | NA | NA | NA | 5.00 | -45.14 | 101.04 |
| NA | -0.15 | NA | 16.89 | 0.71 | NA | NA | NA | -14.74 | 5.00 | -45.15 | 101.04 |
| + | -0.38 | 0.54 | -5.42 | NA | + | 1.36 | -5.88 | NA | 8.00 | -41.59 | 101.06 |
| + | NA | 0.07 | 0.89 | NA | + | NA | NA | NA | 5.00 | -45.17 | 101.08 |
| + | NA | NA | 0.92 | 0.02 | + | NA | NA | NA | 5.00 | -45.17 | 101.09 |
| NA | NA | 0.06 | 16.70 | 0.69 | NA | NA | NA | -14.60 | 5.00 | -45.18 | 101.11 |
| + | -0.24 | NA | 16.71 | 0.64 | NA | NA | NA | -14.38 | 6.00 | -44.04 | 101.15 |
| + | NA | 0.76 | 7.18 | 0.91 | NA | NA | -3.10 | -7.87 | 7.00 | -42.99 | 101.41 |
| NA | -0.12 | NA | 1.49 | NA | NA | 0.78 | NA | NA | 4.00 | -46.56 | 101.62 |
| + | -0.12 | NA | 1.74 | 0.17 | NA | NA | NA | NA | 5.00 | -45.47 | 101.68 |
| + | NA | 0.14 | 1.67 | 0.18 | NA | NA | NA | NA | 5.00 | -45.47 | 101.69 |
| + | -0.09 | 0.08 | 1.67 | NA | NA | NA | NA | NA | 5.00 | -45.49 | 101.74 |
| + | -0.34 | 0.77 | -1.85 | 0.79 | NA | NA | -4.64 | NA | 7.00 | -43.18 | 101.80 |
| + | NA | NA | NA | NA | NA | NA | NA | NA | 2.00 | -48.88 | 101.90 |

| | | | | | | | | | | | |
|---|---|---|---|---|---|---|---|---|---|---|---|
| NA | NA | NA | NA | NA | NA | NA | NA | NA | 1.00 | -49.94 | 101.93 |
| + | -0.28 | NA | 15.62 | 0.61 | + | NA | NA | -14.50 | 7.00 | -43.38 | 102.19 |
| NA | -0.14 | 0.39 | -1.79 | 0.63 | NA | NA | -4.52 | NA | 6.00 | -44.57 | 102.20 |
| + | -0.28 | 0.69 | -4.04 | 0.43 | + | NA | -5.40 | NA | 8.00 | -42.17 | 102.21 |
| + | -0.33 | NA | 0.79 | NA | + | 0.96 | NA | NA | 6.00 | -44.58 | 102.21 |
| NA | NA | -0.23 | 1.68 | 0.08 | NA | NA | NA | NA | 4.00 | -46.87 | 102.23 |
| NA | 0.03 | -0.26 | 1.66 | NA | NA | NA | NA | NA | 4.00 | -46.88 | 102.25 |
| + | NA | 0.25 | 15.63 | 0.63 | + | NA | NA | -14.52 | 7.00 | -43.42 | 102.29 |
| + | -0.68 | 1.06 | 8.24 | 1.21 | + | 0.98 | -6.16 | -14.62 | 10.00 | -39.68 | 102.30 |
| NA | -0.01 | NA | 1.60 | 0.16 | NA | NA | NA | NA | 4.00 | -46.94 | 102.37 |
| + | -0.25 | 0.42 | -0.83 | NA | NA | 0.66 | -3.13 | NA | 7.00 | -43.57 | 102.58 |
| NA | -0.10 | 0.12 | -1.04 | NA | NA | 0.59 | -3.35 | NA | 6.00 | -44.80 | 102.66 |
| NA | -0.29 | NA | 15.88 | 0.75 | NA | 0.78 | NA | -13.84 | 6.00 | -44.81 | 102.69 |
| + | -0.39 | NA | 15.10 | 0.68 | NA | 0.86 | NA | -12.91 | 7.00 | -43.65 | 102.73 |
| + | -0.32 | NA | 1.68 | 0.23 | NA | 0.94 | NA | NA | 6.00 | -44.89 | 102.84 |
| + | -0.52 | 0.79 | -5.60 | 0.61 | + | 1.53 | -6.46 | NA | 9.00 | -41.25 | 102.87 |
| + | NA | 0.63 | NA | NA | NA | NA | NA | NA | 3.00 | -48.29 | 102.88 |
| + | -0.28 | 0.13 | 1.59 | NA | NA | 0.92 | NA | NA | 6.00 | -44.94 | 102.93 |
| + | -0.31 | 0.47 | 18.41 | 0.84 | NA | NA | NA | -16.16 | 7.00 | -43.81 | 103.07 |
| NA | -0.28 | 0.61 | 8.82 | 1.06 | NA | NA | -3.21 | -9.66 | 7.00 | -43.86 | 103.15 |
| + | -0.14 | 0.08 | 0.88 | NA | + | NA | NA | NA | 6.00 | -45.14 | 103.34 |
| NA | -0.16 | 0.08 | 17.13 | 0.75 | NA | NA | NA | -15.00 | 6.00 | -45.14 | 103.34 |
| + | -0.14 | NA | 0.93 | 0.05 | + | NA | NA | NA | 6.00 | -45.14 | 103.34 |
| + | -0.42 | 0.90 | 7.89 | 1.11 | NA | NA | -3.35 | -8.73 | 8.00 | -42.74 | 103.35 |
| + | NA | 0.08 | 0.91 | 0.05 | + | NA | NA | NA | 6.00 | -45.16 | 103.39 |
| + | -0.49 | 0.84 | -1.76 | 0.92 | NA | 0.85 | -4.32 | NA | 8.00 | -42.82 | 103.52 |
| + | -0.46 | NA | 14.64 | 0.65 | + | 0.98 | NA | -13.69 | 8.00 | -42.90 | 103.68 |
| NA | -0.13 | -0.25 | 1.60 | NA | NA | 0.79 | NA | NA | 5.00 | -46.47 | 103.69 |
| NA | -0.17 | NA | 1.55 | 0.21 | NA | 0.82 | NA | NA | 5.00 | -46.50 | 103.76 |
| NA | NA | 0.28 | NA | NA | NA | NA | NA | NA | 2.00 | -49.81 | 103.77 |
| NA | -0.29 | 0.47 | -1.82 | 0.77 | NA | 0.80 | -4.40 | NA | 7.00 | -44.23 | 103.89 |
| + | -0.14 | 0.16 | 1.69 | 0.22 | NA | NA | NA | NA | 6.00 | -45.44 | 103.94 |
| NA | 0.15 | NA | NA | NA | NA | NA | NA | NA | 2.00 | -49.90 | 103.95 |

| | | | | | | | | | | | |
|---|---|---|---|---|---|---|---|---|---|---|---|
| NA | NA | NA | NA | -0.14 | NA | NA | NA | NA | 2.00 | -49.91 | 103.96 |
| + | NA | NA | NA | -0.10 | NA | NA | NA | NA | 3.00 | -48.86 | 104.02 |
| + | 0.08 | NA | NA | NA | NA | NA | NA | NA | 3.00 | -48.87 | 104.03 |
| + | -0.32 | 0.30 | 16.40 | 0.74 | + | NA | NA | -15.25 | 8.00 | -43.28 | 104.43 |
| + | -0.50 | 0.60 | 17.36 | 0.96 | NA | 0.98 | NA | -15.30 | 8.00 | -43.30 | 104.47 |
| NA | 0.01 | -0.23 | 1.67 | 0.08 | NA | NA | NA | NA | 5.00 | -46.87 | 104.49 |
| + | -0.34 | 0.11 | 0.75 | NA | + | 0.96 | NA | NA | 7.00 | -44.56 | 104.56 |
| + | -0.35 | NA | 0.85 | 0.10 | + | 0.97 | NA | NA | 7.00 | -44.56 | 104.56 |
| NA | -0.31 | 0.16 | 16.44 | 0.83 | NA | 0.81 | NA | -14.43 | 7.00 | -44.78 | 105.00 |
| + | NA | 0.67 | NA | 0.11 | NA | NA | NA | NA | 4.00 | -48.28 | 105.05 |
| + | 0.02 | 0.63 | NA | NA | NA | NA | NA | NA | 4.00 | -48.29 | 105.08 |
| + | -0.35 | 0.23 | 1.60 | 0.30 | NA | 0.96 | NA | NA | 7.00 | -44.83 | 105.09 |
| NA | -0.36 | 0.63 | 8.06 | 1.09 | NA | 0.56 | -3.04 | -8.88 | 8.00 | -43.72 | 105.31 |
| + | -0.53 | 0.94 | 7.45 | 1.16 | NA | 0.74 | -2.99 | -8.15 | 9.00 | -42.50 | 105.38 |
| + | -0.15 | 0.11 | 0.92 | 0.09 | + | NA | NA | NA | 7.00 | -45.13 | 105.69 |
| + | -0.53 | 0.43 | 15.81 | 0.84 | + | 1.06 | NA | -14.85 | 9.00 | -42.72 | 105.81 |
| NA | 0.14 | 0.28 | NA | NA | NA | NA | NA | NA | 3.00 | -49.78 | 105.85 |
| NA | NA | 0.26 | NA | -0.05 | NA | NA | NA | NA | 3.00 | -49.81 | 105.91 |
| NA | -0.16 | -0.21 | 1.62 | 0.14 | NA | 0.81 | NA | NA | 6.00 | -46.45 | 105.96 |
| NA | 0.18 | NA | NA | -0.18 | NA | NA | NA | NA | 3.00 | -49.85 | 106.00 |
| + | 0.10 | NA | NA | -0.12 | NA | NA | NA | NA | 4.00 | -48.85 | 106.19 |
| + | -0.37 | 0.16 | 0.82 | 0.16 | + | 0.98 | NA | NA | 8.00 | -44.53 | 106.94 |
| + | 0.00 | 0.67 | NA | 0.11 | NA | NA | NA | NA | 5.00 | -48.28 | 107.31 |
| NA | 0.16 | 0.24 | NA | -0.09 | NA | NA | NA | NA | 4.00 | -49.77 | 108.02 |

**Table S5.** Statistical results for endophyte prevalence modeled as a beta regression of historical endophyte prevalence, mean SPEI, and SPEI SD. This was determined to be the best-supported model through global model selection.

| | $\chi^2$ | DF | P |
|---|---|---|---|
| Historical endophyte prevalence | 27.35 | 1 | < 0.0001 |
| Mean SPEI | 3.31 | 1 | 0.069 |
| SPEI SD | 10.52 | 1 | 0.0012 |

**Table S6.** Comparison of candidate models for endophyte prevalence.

| **Fire occurrenc** | **Mean SPEI** | **Coefficient of** | **Historical endophyte** | **SPEI SD** | **Endophyte :fire** | **Endophyte :mean** | **Endophyte :SPEI** | **Endophyte :SPEI SD** | **DF** | **Log-likelihood** | **AICc** |
|---|---|---|---|---|---|---|---|---|---|---|---|

| e | | regression of SPEI on time | prevalence | | | SPEI | coefficient of regression | | | | |
|---|---|---|---|---|---|---|---|---|---|---|---|
| NA | 1.26 | NA | 3.36 | -2.58 | NA | NA | NA | NA | 6.00 | -31.76 | 77.03 |
| NA | NA | NA | 3.54 | -2.34 | NA | NA | NA | NA | 5.00 | -33.54 | 78.12 |
| NA | 1.36 | -0.88 | 3.59 | -2.90 | NA | NA | NA | NA | 7.00 | -31.22 | 78.47 |
| NA | 1.95 | NA | 3.59 | -2.67 | NA | -0.63 | NA | NA | 7.00 | -31.33 | 78.70 |
| NA | 1.15 | NA | 9.72 | -1.62 | NA | NA | NA | -6.08 | 7.00 | -31.45 | 78.94 |
| NA | NA | NA | 12.49 | -1.00 | NA | NA | NA | -8.60 | 6.00 | -32.89 | 79.28 |
| + | 1.24 | NA | 3.35 | -2.58 | NA | NA | NA | NA | 7.00 | -31.76 | 79.55 |
| NA | NA | -0.60 | 3.72 | -2.53 | NA | NA | NA | NA | 6.00 | -33.29 | 80.07 |
| NA | 2.01 | -0.88 | 3.84 | -3.01 | NA | -0.62 | NA | NA | 8.00 | -30.78 | 80.23 |
| + | NA | NA | 3.49 | -2.41 | NA | NA | NA | NA | 6.00 | -33.44 | 80.37 |
| + | 1.31 | -1.36 | 3.60 | -3.08 | NA | NA | NA | NA | 8.00 | -30.85 | 80.38 |
| NA | 1.23 | -0.58 | 8.32 | -2.07 | NA | NA | NA | -4.59 | 8.00 | -31.08 | 80.84 |
| NA | 1.74 | NA | 8.70 | -1.89 | NA | -0.54 | NA | -4.91 | 8.00 | -31.14 | 80.94 |
| NA | 1.36 | -0.65 | 3.16 | -2.86 | NA | NA | -0.40 | NA | 8.00 | -31.18 | 81.02 |
| + | 1.93 | NA | 3.57 | -2.67 | NA | -0.63 | NA | NA | 8.00 | -31.32 | 81.32 |
| NA | NA | -0.42 | 12.08 | -1.19 | NA | NA | NA | -8.08 | 7.00 | -32.74 | 81.52 |
| + | 1.13 | NA | 9.74 | -1.61 | NA | NA | NA | -6.11 | 8.00 | -31.44 | 81.55 |
| + | NA | NA | 12.40 | -1.04 | NA | NA | NA | -8.57 | 7.00 | -32.80 | 81.65 |
| + | NA | -1.15 | 3.75 | -2.83 | NA | NA | NA | NA | 7.00 | -32.82 | 81.68 |
| + | 1.25 | NA | 3.22 | -2.59 | + | NA | NA | NA | 8.00 | -31.74 | 82.14 |
| + | 1.97 | -1.43 | 3.86 | -3.21 | NA | -0.65 | NA | NA | 9.00 | -30.38 | 82.15 |
| NA | NA | -0.70 | 3.62 | -2.58 | NA | NA | -0.15 | NA | 7.00 | -33.29 | 82.62 |
| + | NA | NA | 3.40 | -2.42 | + | NA | NA | NA | 7.00 | -33.43 | 82.89 |
| NA | 1.82 | -0.57 | 7.40 | -2.33 | NA | -0.55 | NA | -3.51 | 9.00 | -30.75 | 82.90 |
| NA | 2.02 | -0.76 | 3.53 | -3.01 | NA | -0.62 | -0.30 | NA | 9.00 | -30.76 | 82.91 |
| + | 1.21 | -1.07 | 7.49 | -2.35 | NA | NA | NA | -3.77 | 9.00 | -30.77 | 82.94 |
| + | 1.32 | -1.37 | 3.44 | -3.10 | + | NA | NA | NA | 9.00 | -30.81 | 83.02 |
| + | 1.31 | -1.23 | 3.21 | -3.09 | NA | NA | -0.37 | NA | 9.00 | -30.81 | 83.02 |
| NA | 1.20 | -0.22 | 10.26 | -1.60 | NA | NA | -1.00 | -7.43 | 9.00 | -30.86 | 83.11 |
| NA | NA | 0.00 | 12.28 | -1.06 | NA | NA | -1.10 | -9.36 | 8.00 | -32.38 | 83.43 |
| + | NA | -0.87 | 11.02 | -1.54 | NA | NA | NA | -7.04 | 8.00 | -32.40 | 83.46 |
| + | 1.72 | NA | 8.84 | -1.86 | NA | -0.55 | NA | -5.08 | 9.00 | -31.12 | 83.64 |

| | | | | | | | | | | | |
|---|---|---|---|---|---|---|---|---|---|---|---|
| + | 1.97 | NA | 3.39 | -2.70 | + | -0.66 | NA | NA | 9.00 | -31.28 | 83.95 |
| + | NA | -0.90 | 3.21 | -2.63 | NA | NA | -0.47 | NA | 8.00 | -32.73 | 84.13 |
| + | NA | NA | 12.35 | -1.04 | + | NA | NA | -8.65 | 8.00 | -32.79 | 84.24 |
| + | 1.13 | NA | 9.57 | -1.63 | + | NA | NA | -6.07 | 9.00 | -31.42 | 84.25 |
| + | NA | -1.23 | 3.65 | -2.88 | + | NA | NA | NA | 8.00 | -32.82 | 84.30 |
| + | 1.99 | -1.36 | 3.63 | -3.21 | + | -0.67 | NA | NA | 10.00 | -30.31 | 84.85 |
| + | 1.97 | -1.23 | 3.48 | -3.18 | NA | -0.64 | -0.35 | NA | 10.00 | -30.35 | 84.93 |
| + | 1.83 | -1.07 | 6.61 | -2.62 | NA | -0.59 | NA | -2.72 | 10.00 | -30.38 | 84.99 |
| + | 1.17 | -0.69 | 8.63 | -2.04 | NA | NA | -1.02 | -5.89 | 10.00 | -30.55 | 85.34 |
| NA | 1.77 | -0.16 | 8.82 | -1.93 | NA | -0.52 | -1.03 | -5.89 | 10.00 | -30.55 | 85.34 |
| + | NA | -0.44 | 12.40 | -1.16 | NA | NA | -1.24 | -9.61 | 9.00 | -31.98 | 85.35 |
| + | 1.34 | -1.04 | 2.54 | -3.09 | + | NA | -0.73 | NA | 10.00 | -30.70 | 85.64 |
| + | 1.21 | -1.02 | 7.30 | -2.35 | + | NA | NA | -3.73 | 10.00 | -30.75 | 85.74 |
| + | NA | -0.83 | 11.04 | -1.51 | + | NA | NA | -7.19 | 9.00 | -32.38 | 86.15 |
| + | 1.74 | NA | 8.49 | -1.91 | + | -0.57 | NA | -4.90 | 10.00 | -31.08 | 86.40 |
| + | NA | -0.46 | 2.26 | -2.79 | + | NA | -1.10 | NA | 9.00 | -32.58 | 86.56 |
| NA | NA | NA | 3.60 | NA | NA | NA | NA | NA | 4.00 | -39.28 | 87.26 |
| + | NA | 0.02 | 12.16 | -1.00 | + | NA | -2.00 | -10.63 | 10.00 | -31.67 | 87.56 |
| + | 2.03 | -1.12 | 2.80 | -3.22 | + | -0.68 | -0.68 | NA | 11.00 | -30.21 | 87.59 |
| + | 1.75 | -0.67 | 7.98 | -2.24 | NA | -0.55 | -0.99 | -5.04 | 11.00 | -30.22 | 87.61 |
| + | 1.14 | -0.21 | 9.06 | -1.72 | + | NA | -1.78 | -7.51 | 11.00 | -30.32 | 87.82 |
| + | 1.85 | -1.02 | 6.18 | -2.67 | + | -0.61 | NA | -2.50 | 11.00 | -30.34 | 87.86 |
| NA | 0.66 | NA | 3.54 | NA | NA | NA | NA | NA | 5.00 | -38.70 | 88.45 |
| + | NA | NA | 3.65 | NA | NA | NA | NA | NA | 5.00 | -39.22 | 89.50 |
| NA | NA | 0.17 | 3.54 | NA | NA | NA | NA | NA | 5.00 | -39.25 | 89.54 |
| + | 1.76 | -0.18 | 8.22 | -1.97 | + | -0.58 | -1.80 | -6.56 | 12.00 | -29.97 | 90.19 |
| NA | 0.96 | NA | 3.71 | NA | NA | -0.35 | NA | NA | 6.00 | -38.54 | 90.58 |
| + | 0.68 | NA | 3.61 | NA | NA | NA | NA | NA | 6.00 | -38.61 | 90.73 |
| NA | 0.66 | 0.04 | 3.53 | NA | NA | NA | NA | NA | 6.00 | -38.68 | 90.86 |
| + | NA | 0.37 | 3.57 | NA | NA | NA | NA | NA | 6.00 | -39.10 | 91.71 |
| + | NA | NA | 3.86 | NA | + | NA | NA | NA | 6.00 | -39.16 | 91.82 |
| NA | NA | 0.37 | 3.00 | NA | NA | NA | -0.51 | NA | 6.00 | -39.19 | 91.87 |
| + | 0.74 | 0.45 | 3.50 | NA | NA | NA | NA | NA | 7.00 | -38.43 | 92.90 |

| | | | | | | | | | | | |
|---|---|---|---|---|---|---|---|---|---|---|---|
| NA | 1.06 | 0.30 | 3.63 | NA | NA | -0.42 | NA | NA | 7.00 | -38.44 | 92.91 |
| + | 0.95 | NA | 3.76 | NA | NA | -0.33 | NA | NA | 7.00 | -38.47 | 92.97 |
| + | 0.70 | NA | 3.86 | NA | + | NA | NA | NA | 7.00 | -38.52 | 93.08 |
| NA | 0.72 | 0.27 | 2.94 | NA | NA | NA | -0.54 | NA | 7.00 | -38.54 | 93.11 |
| + | NA | 0.33 | 3.75 | NA | + | NA | NA | NA | 7.00 | -39.07 | 94.17 |
| + | NA | 0.36 | 3.28 | NA | NA | NA | -0.29 | NA | 7.00 | -39.07 | 94.18 |
| + | 1.03 | 0.26 | 3.71 | NA | NA | -0.38 | NA | NA | 8.00 | -38.28 | 95.23 |
| NA | 1.15 | 0.60 | 2.77 | NA | NA | -0.41 | -0.81 | NA | 8.00 | -38.29 | 95.26 |
| + | 0.81 | 0.74 | 2.65 | NA | NA | NA | -0.81 | NA | 8.00 | -38.30 | 95.26 |
| + | 1.00 | NA | 4.03 | NA | + | -0.35 | NA | NA | 8.00 | -38.37 | 95.41 |
| + | 0.75 | 0.40 | 3.70 | NA | + | NA | NA | NA | 8.00 | -38.38 | 95.44 |
| + | NA | 0.27 | 3.60 | NA | + | NA | -0.15 | NA | 8.00 | -39.06 | 96.79 |
| + | 1.13 | 0.48 | 3.23 | NA | NA | -0.40 | -0.44 | NA | 9.00 | -38.15 | 97.69 |
| + | 1.04 | 0.21 | 3.95 | NA | + | -0.38 | NA | NA | 9.00 | -38.24 | 97.87 |
| + | 0.81 | 0.69 | 2.80 | NA | + | NA | -0.73 | NA | 9.00 | -38.29 | 97.98 |
| + | 1.09 | 0.39 | 3.58 | NA | + | -0.39 | -0.28 | NA | 10.00 | -38.15 | 100.53 |
| NA | 1.85 | NA | NA | -2.93 | NA | NA | NA | NA | 5.00 | -47.73 | 106.52 |
| + | 1.76 | NA | NA | -3.04 | NA | NA | NA | NA | 6.00 | -46.92 | 107.34 |
| NA | 1.81 | 0.58 | NA | -2.71 | NA | NA | NA | NA | 6.00 | -47.45 | 108.39 |
| + | 1.76 | 0.03 | NA | -3.03 | NA | NA | NA | NA | 7.00 | -46.92 | 109.87 |
| + | NA | NA | NA | -2.68 | NA | NA | NA | NA | 5.00 | -51.03 | 113.10 |
| NA | NA | NA | NA | -2.41 | NA | NA | NA | NA | 4.00 | -52.29 | 113.27 |
| NA | NA | 0.81 | NA | -2.22 | NA | NA | NA | NA | 5.00 | -51.58 | 114.21 |
| + | NA | 0.35 | NA | -2.56 | NA | NA | NA | NA | 6.00 | -50.93 | 115.35 |
| NA | 1.09 | 1.36 | NA | NA | NA | NA | NA | NA | 5.00 | -56.16 | 123.37 |
| NA | NA | 1.26 | NA | NA | NA | NA | NA | NA | 4.00 | -57.97 | 124.63 |
| + | 1.16 | 1.60 | NA | NA | NA | NA | NA | NA | 6.00 | -55.94 | 125.37 |
| + | NA | 1.40 | NA | NA | NA | NA | NA | NA | 5.00 | -57.88 | 126.82 |
| NA | 0.87 | NA | NA | NA | NA | NA | NA | NA | 4.00 | -59.19 | 127.07 |
| NA | NA | NA | NA | NA | NA | NA | NA | NA | 3.00 | -60.54 | 127.48 |
| + | 0.86 | NA | NA | NA | NA | NA | NA | NA | 5.00 | -58.92 | 128.90 |
| + | NA | NA | NA | NA | NA | NA | NA | NA | 4.00 | -60.23 | 129.16 |

**Table S7.** Information on common garden sites used for population modeling.

| **Latitude** | **Longitude** | **Year 1 SPEI** | **Year 2 SPEI** | **Year 3 SPEI** | **Year 4 SPEI** |
|---|---|---|---|---|---|
| 39.74 | -123.63 | 0.86 | 1.39 | 0.92 | 0.40 |
| 36.38 | -121.57 | 0.15 | 0.44 | 0.15 | 0.08 |
| 38.87 | -122.45 | -0.32 | 0.38 | -0.12 | -0.55 |
| 38.49 | -122.15 | -0.70 | -0.13 | -0.44 | -0.78 |
| 38.86 | -122.42 | -0.44 | 0.26 | -0.21 | -0.65 |

**Table S8.** Information on source populations used in the common garden experiments for population modeling.

| **Latitude** | **Longitude** | **Endophyte prevalence** | **Endophyte status** |
|---|---|---|---|
| 38.48 | -122.15 | 0 | - |
| 39.15 | -122.75 | 0 | - |
| 39.74 | -123.63 | 0 | - |
| 39.05 | -122.49 | 0 | - |
| 39.41 | -122.60 | 0 | - |
| 39.46 | -120.87 | 0 | - |
| 39.10 | -120.92 | 25 | + |
| 39.34 | -122.70 | 76.7 | + |
| 39.20 | -123.29 | 76.9 | + |
| 36.38 | -121.57 | 90 | + |
| 39.12 | -122.81 | 100 | + |

**Table S9.** Vital rate functions and terms in their respective selected models.

| Vital rate | Fixed effects |
|---|---|
| **Survival** | 0.77*size + 1.51*SPEI – 0.02*damage + 0.09*endophyte – 0.08*size$^2$ – 0.19*size:endophyte + 0.53*SPEI:endophyte – 0.19*damage:endophyte + 0.31*age1 – 0.38*age2 – 0.22*age3 + 0.25*age4 + (1\|population) + (1\|garden) + 0.55 |
| Growth | 0.39*size + 0.04*damage – 0.03*size$^2$ + 0.27*age1 + 0.16*age2 + 0.22*age3 + 0.28*age4 + (1\|population) + (1\|garden) + 0.39 |
| **Growth standard deviation** | 0.37*size + 0.06*endophyte – 0.17*SPEI + 0.02*fungicide + 0.11*damage – 0.03*size$^2$ + 0.067*age1 – 0.0007*age2 + 0.01*age3 + 0.20*age4 + 0.18*SPEI:fungicide + 0.17*endophyte:SPEI – 0.15*endophyte:fungicide – 0.07*endophyte:damage – 0.15*endophyte:SPEI:fungicide + 0.03 |
| **Flowering probability** | 0.98*size + 0.15*SPEI + 0.55*endophyte + 0.29*fungicide – 0.10*size$^2$ – 3.59*age1 + 0.03*age2 – 0.82*age3 + 0.29*age4 + 0.22*age6 – 0.50*SPEI:endophyte – 0.63*SPEI:fungicide – 0.67*endophyte:fungicide + 0.84*SPEI:endophyte:fungicide + (1\|population) + (1\|garden) – 1.07 |
| **Flower number** | 0.08*size – 0.09*endophyte – 0.77*age1 – 0.62*age2 – 0.52*age3 – 0.15*age4 – 0.08*age6 + 0.08*size:endophyte + (1\|population) + (1\|garden)+ 0.63 |
| Seeds per tiller | parameter; 11.06 |
| First year | 1.24 + (1\|population) + (1\|garden) |

| | |
|---|---|
| germination probability | |
| **Size of germinants** | 0.29*endophyte + 0.09*fungicide – 0.35*endophyte:fungicide + (1\|population) + (1\|garden) + 0.23 |
| Size of germinants standard deviation | 0.77 |
| **Probability of entering dormancy** | 0.16*size + 0.25*damage – 0.58*endophyte – 0.17*fungicide – 1.58*age1 – 0.45*age2 – 1.01*age3 – 0.49*size:endophyte – 0.67*damage:endophyte – 1.52*damage:fungicide + (1\|population) + (1\|garden) – 2.79 |
| Probability of leaving dormancy | 0.32 + (1\|population) + (1\|garden) |
| **Size out of dormancy** | -0.51*SPEI + 0.10*endophyte + 0.07*fungicide + (1\|population) + 0.63 |
| Size out of dormancy standard deviation | 1.82 |
| Probability of seed survival in seedbank | parameter; 0.34 |
| Probability of germination from seedbank | parameter; 0.21 |

NOTE: Bolded vital rates are those involving endophyte status. SPEI=SPEI of a given site the year of sampling, damage=damage per tiller, endophyte=endophyte status (associated, free). Age also corresponds to year of experiment. Colons (:) indicate interaction terms. The random effects (population of origin and common garden site) were excluded from the growth standard deviation model to allow for model convergence.

**Table S10.** Fit of vital rate functions including the plant individual random effect in comparison to those excluding the individual random effect.

| | $\chi^2$ | DF | P |
|---|---|---|---|
| Survival | 0.053 | 1 | 0.82 |
| Growth | ~0 | 1 | ~1 |
| Flowering probability | 75.91 | 1 | < 0.0001 |
| Flower number | ~0 | 1 | ~1 |
| Probability of entering dormancy | 0.27 | 1 | 0.60 |

**Table S11.** Statistical results for the difference in modeled λ between hypothetical endophyte-associated and endophyte-free populations as a function of mean site SPEI.

| | F | DF | P |
|---|---|---|---|
| Mean site $SPEI^2$ | 584.31 | 2, 83 | < 0.0001 |

**Table S12.** Statistical results for endophyte contribution to population growth rate (i.e., the difference in λ between modeled endophyte-associated and endophyte-free populations for a range of SPEI conditions across the host range) as a function of site SPEI SD.

| | F | DF | P |
|---|---|---|---|
| Site SPEI SD | 3.68 | 1, 84 | 0.058 |

**Table S13.** Statistical results for a linear regression between historical endophyte prevalence in our original 2009 surveys and pre-survey interannual variability in aridity (i.e., SPEI SD) between 1995–2008.

| | F | DF | P |
|---|---|---|---|
| Historical SPEI SD | 22.34 | 1, 61 | < 0.0001 |
